# Interleukin 11 therapy causes acute heart failure and its use in patients should be reconsidered

**DOI:** 10.1101/2023.09.30.560259

**Authors:** Mark Sweeney, Katie O’Fee, Chelsie Villanueva-Hayes, Ekhlas Rahman, Michael Lee, Henrike Maatz, Eric L. Lindberg, Konstantinos Vanezis, Ivan Andrew, Emma R. Jennings, Wei-Wen Lim, Anissa A Widjaja, Norbert Hubner, Paul J.R. Barton, Stuart A Cook

## Abstract

**Background:** Interleukin 11 (IL11) was initially thought important for platelet production, which led to recombinant IL11 being developed as a drug to treat thrombocytopenia. IL11 was later found to be redundant for haematopoiesis and its use in patients is associated with unexplained cardiac side effects. Here we identify previously unappreciated and direct cardiomyocyte toxicities associated with IL11 therapy.

**Methods:** We injected recombinant mouse lL11 (rmIL11) into mice and studied its molecular effects in the heart using immunoblotting, qRT-PCR, bulk RNA-seq, single nuclei RNA-seq (snRNA-seq) and ATAC-seq. The physiological impact of IL11 was assessed by echocardiography *in vivo* and using cardiomyocyte contractility assays *in vitro*. To determine the activity of IL11 specifically in cardiomyocytes we made two cardiomyocyte-specific *Il11ra1* knockout mouse models using either AAV9-mediated and *Tnnt2*-restricted (vCMKO) or *Myh6* (m6CMKO) Cre expression and an *Il11ra1* floxed mouse strain. In pharmacologic studies, we studied the effects of JAK/STAT inhibition on rmIL11-induced cardiac toxicities.

**Results:** Injection of rmIL11 caused acute and dose-dependent impairment of left ventricular ejection fraction (saline (2 µL/kg), 60.4%±3.1; rmIL11 (200 mcg/kg), 31.6%±2.0; p<0.0001, n=5). Following rmIL11 injection, myocardial STAT3 and JNK phosphorylation were increased and bulk RNA-seq revealed upregulation of pro-inflammatory pathways (TNFα, NFκB and JAK/STAT) and perturbed calcium handling. SnRNA-seq showed rmIL11-induced expression of stress factors (*Ankrd1*, *Ankrd23*, *Xirp2*), activator protein-1 (AP-1) transcription factor genes and *Nppb* in the cardiomyocyte compartment. Following rmIL11 injection, ATAC-seq identified epigenetic enrichment of the *Ankrd1* and *Nppb* genes and stress-responsive, AP-1 transcription factor binding sites. Cardiomyocyte-specific effects were examined in vCMKO and m6CMKO mice, which were both protected from rmIL11-induced left ventricular impairment and molecular pathobiologies. In mechanistic studies, inhibition of JAK/STAT signalling with either ruxolitinib or tofacitinib prevented rmIL11-induced cardiac dysfunction.

**Conclusions:** Injection of IL11 directly activates JAK/STAT3 in cardiomyocytes to cause acute heart failure. Our data overturn the earlier assumption that IL11 is cardioprotective and explain the serious cardiac side effects associated with IL11 therapy, which questions its continued use in patients.

**Clinical Perspective:** *What is new?:*

- Injection of IL11 to mice causes acute and dose-dependent left ventricular impairment
- IL11 activates JAK/STAT3 in cardiomyocytes to cause cell stress, inflammation and impaired calcium handling
- These data identify, for the first time, that IL11 is directly toxic in cardiomyocytes, overturning the earlier literature that suggested the opposite

*What are the clinical implications?:*

- Recombinant human IL11 (rhIL11) is used as a drug to increase platelets in patients with thrombocytopenia but this has severe and unexplained cardiac side effects
- We show that IL11 injection causes cardiomyocyte dysfunction and heart failure, which explains its cardiac toxicities that were previously thought non-specific
- These findings have immediate translational implications as they question the continued use of rhIL11 in patients around the world

## Introduction

Interleukin 11 (IL11) is an elusive member of the interleukin 6 (IL6) family of cytokines, which collectively signal via the gp130 co-receptor. Following its identification in 1990^1^ recombinant human IL11 (rhIL11) was found to increase megakaryocyte activity and peripheral platelet counts in mice^2^. Soon after, IL11 was developed as a therapeutic (Oprelvekin; Neumega) to increase platelet counts in patients with chemotherapy-induced thrombocytopenia, received FDA approval for this indication in 1998 and is still used to this day^3,4^. In recent years, longer-acting formulations of rhIL11 have been tested in pre-clinical studies and new clinical trials of PEGylated rhIL11 in patients are anticipated^5^.

RhIL11 was also trialled to increase platelet counts in patients with von Willebrand factor deficiency, myelodysplastic syndrome, cirrhosis and sepsis, and tested as a putative cytoprotective agent in numerous other conditions, including myocardial infarction^6^ [**Table 1 and Suppl Table 1**]. However, it became apparent that IL11 is not required for basal or compensatory blood cell or platelet counts in mice or humans: IL11 is in fact redundant for haematopoiesis^7,8^. Thus, the effects of injection of high dose rhIL11 on platelets appear non-physiological and possibly reflect non-specific gp130 activity^9,10^.

**Table 1.**
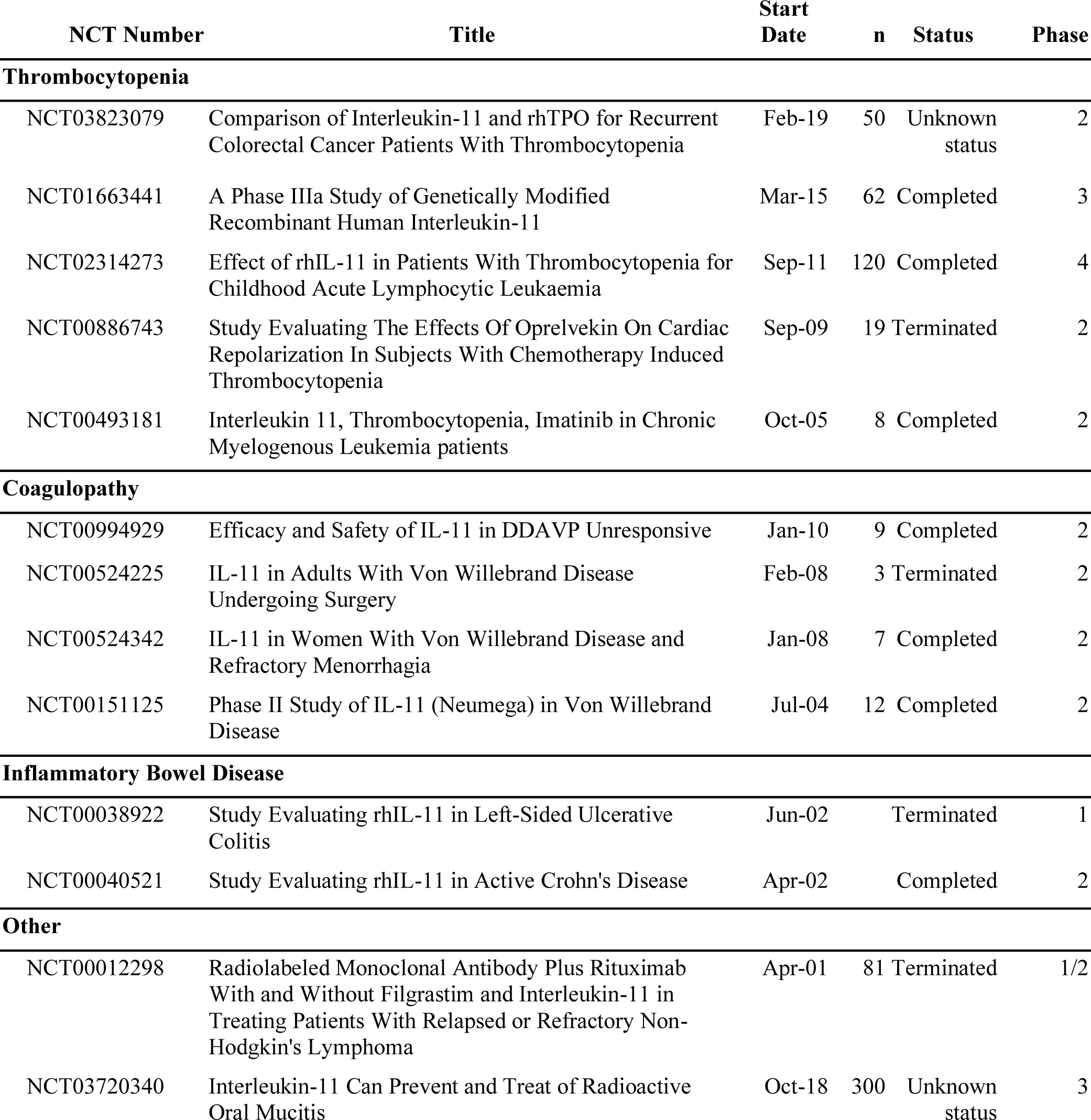
Human clinical trials registered with clinicaltrials.gov using recombinant human interleukin 11.

| NCT Number | Title | Start Date | n | Status | Phase |
| --- | --- | --- | --- | --- | --- |
| **Thrombocytopenia** |  |  |  |  |  |
| NCT03823079 | Comparison of Interleukin-11 and rhTPO for Recurrent Colorectal Cancer Patients With Thrombocytopenia | Feb-19 | 50 | Unknown status | 2 |
| NCT01663441 | A Phase IIIa Study of Genetically Modified Recombinant Human Interleukin-11 | Mar-15 | 62 | Completed | 3 |
| NCT02314273 | Effect of rhIL-11 in Patients With Thrombocytopenia for Childhood Acute Lymphocytic Leukaemia | Sep-11 | 120 | Completed | 4 |
| NCT00886743 | Study Evaluating The Effects Of Oprelvekin On Cardiac Repolarization In Subjects With Chemotherapy Induced Thrombocytopenia | Sep-09 | 19 | Terminated | 2 |
| NCT00493181 | Interleukin 11, Thrombocytopenia, Imatinib in Chronic Myelogenous Leukemia patients | Oct-05 | 8 | Completed | 2 |
| **Coagulopathy** |  |  |  |  |  |
| NCT00994929 | Efficacy and Safety of IL-11 in DDAVP Unresponsive | Jan-10 | 9 | Completed | 2 |
| NCT00524225 | IL-11 in Adults With Von Willebrand Disease Undergoing Surgery | Feb-08 | 3 | Terminated | 2 |
| NCT00524342 | IL-11 in Women With Von Willebrand Disease and Refractory Menorrhagia | Jan-08 | 7 | Completed | 2 |
| NCT00151125 | Phase II Study of IL-11 (Neumega) in Von Willebrand Disease | Jul-04 | 12 | Completed | 2 |
| **Inflammatory Bowel Disease** |  |  |  |  |  |
| NCT00038922 | Study Evaluating rhIL-11 in Left-Sided Ulcerative Colitis | Jun-02 |  | Terminated | 1 |
| NCT00040521 | Study Evaluating rhIL-11 in Active Crohn's Disease | Apr-02 |  | Completed | 2 |
| **Other** |  |  |  |  |  |
| NCT00012298 | Radiolabeled Monoclonal Antibody Plus Rituximab With and Without Filgrastim and Interleukin-11 in Treating Patients With Relapsed or Refractory Non-Hodgkin's Lymphoma | Apr-01 | 81 | Terminated | 1/2 |
| NCT03720340 | Interleukin-11 Can Prevent and Treat of Radioactive Oral Mucitis | Oct-18 | 300 | Unknown status | 3 |

Unfortunately, injection of rhIL11 into patients has severe and hitherto unexplained cardiac side effects. Up to 20% of patients given rhIL11 (50 mcg/kg) develop atrial arrhythmias, a high proportion of individuals develop heart failure and rare cases of ventricular arrhythmias and sudden death are reported^11,12^. Furthermore, serum natriuretic peptide levels become acutely and transiently elevated in patients receiving IL11 therapy, with B-natriuretic peptide levels sometimes exceeding those diagnostic of heart failure.

While IL11 was previously thought to be cytoprotective, anti-inflammatory and anti-fibrotic in the heart^13–15^ and other organs, recent studies by ourselves and others have challenged this premise^16–18^. Indeed, experiments over the last five years have questioned the earlier literature and IL11 is increasingly viewed as pro-inflammatory and pro-fibrotic. Given this large shift in our understanding of IL11 and the fact that cardiomyocytes (CMs) robustly express IL11RA, we devised experiments to determine whether IL11 is toxic to CMs and if this could explain cardiac side effects associated with IL11 therapy in patients.

## Methods

Detailed information on experimental methods of RNA and DNA analysis and cardiomyocyte isolation protocols are provided in the supplementary material.

### Animal studies

All mouse studies were conducted according to the Animals (Scientific Procedures) Act 1986 Amendment Regulations 2012 and approved by the Animal Welfare Ethical Review Body at Imperial College London. Animal experiments were carried out under UK Home Office Project License P108022D1 (September 2019). Wild type (WT) mice on a C57BL/6J background were purchased from Charles River (Cat#632). They were bred in a dedicated breeding facility and housed in a single room of the experimental animal facility with a 12-hour light-dark cycle and provided food and water *ad libitum*. The *Il11ra1* floxed mouse (C57BL/6-Il11ra1^em1Cook^/J, Jax:034465) has exons 4-7 of the *Il11ra1* gene flanked by loxP sites as has been described previously^19^. In the presence of Cre-recombinase excision of exon 4-7 results in a non-functional IL11 receptor.

Male myosin heavy chain 6 Cre (*Myh6*-Cre) mice (B6.FVB-Tg(Myh6-cre)2182Mds/J, Jax:011038) were purchased from Jax (Bar Harbor, Maine, United States) as heterozygotes. These mice were crossed with homozygous *Il11ra1* floxed females. In the second generation, mice from generation one, heterozygous for the *Il11ra1* flox allele and heterozygous for the Cre, were crossed with *Il11ra1* flox homozygotes to produce littermate experimental and control animals.

Recombinant mouse interleukin-11 (rmIL11) (Z03052, Genscript, Oxford, UK) was dissolved in phosphate-buffered saline (PBS) (14190144, ThermoFisher, MA, USA), and injected intraperitoneally (IP) at a dose of 200 mcg/kg unless otherwise stated. Control mice received an equivalent volume of saline (2 µL/kg). Recombinant mouse interleukin-6 (Z02767, Genscript) was dissolved in PBS and injected IP at a dose of 200 mcg/kg. Mice were randomly assigned to a treatment group using a random number generator and syringes for injection were prepared and blinded by a different investigator than administered the IP injection.

### Genotyping

Genotype was confirmed with ear-notch DNA samples. DNA was extracted using a sodium hydroxide digestion buffer, then neutralised with 1M Tris-HCl pH 8. *Il11ra1* flox genotype was confirmed with a single polymerase chain reaction (PCR) reaction yielding a PCR product at 163bp for the wild type allele or 197bp in the transgenic allele. *Myh6*-Cre mice were genotyped using two reactions for either the transgenic gene product of 295bp (or wild type gene product of 300bp) along with an internal positive control (200bp). Primers used in these reactions are detailed in supplementary table 2.

#### Viral Vector

The viral vector used in this study, AAV9-cTNT-EGFP-T2A-iCre-WPRE (VB5413), was purchased from Vector Biolabs (Malvern, PA, USA). A codon optimised Cre was delivered using an AAV9 capsid and under the control of the *Tnnt2* promoter. This was linked to an enhanced green fluorescent protein (EGFP) reporter with a 2a self-cleaving linker. 1×10^12^ genome copies or an equivalent volume of saline were injected into the tail veins of 8 - 9 week old homozygous male *Il11ra1* flox mice and from this point mice were housed separately from saline-injected controls for 3 weeks prior to further experiments.

### Echocardiography

Echocardiography was performed under light isoflurane anaesthesia using a Vevo3100 imaging system and MX550D linear transducer (Fujifilm Visualsonic Inc, ON, Canada).

Anaesthesia was induced with 4% isoflurane for 1 minute and maintained with 1-2% isoflurane. Mice were allowed to equilibrate to the anaesthetic after induction for 9 minutes before imaging was started. Heart rate measurement from single lead electrocardiogram (ECG) recordings were taken at the completion of the equilibration period. Measurements of ventricular ejection fraction (LVEF) was measured from m-mode images taken in the parasternal short axis (PSAX) view at midventricular level and averaged across 3 heartbeats. Global circumferential strain (GCS) measurements were also taken from the PSAX view and analysed in a semi-automated fashion by the VevoStrain imaging software (VevoLab, version 5.5.0, Fujifilm Visualsonic). Aortic velocity time integral (VTI) was measured using pulse wave doppler in the aortic arch and an average taken from 3 heart beats. The investigator was blinded to the treatment group for all studies at both the imaging acquisition and analysis stages.

### qPCR

The tissue was washed in ice-cold PBS and snap-frozen in liquid nitrogen. Total RNA was extracted using TRIzol (15596026, Invitrogen, MA, USA,) in RNeasy columns (74106, Qiagen, MD, USA). cDNA was synthesised using Superscript Vilo Mastermix (11755050, Invitrogen). Gene expression analysis was performed using quantitative polymerase chain reaction (qPCR) with TaqMan gene expression assay in duplicate over 40 cycles. *Il11ra1*: custom TaqMan assay **[Suppl Table 3]**, *Nppb*: Mm01255770_g1, *Rrad*: Mm00451053_m1. Gene expression data were normalised to *Gapdh* expression (Mm99999915_g1) expression and fold change compared to control samples was calculated using 2^-ΔΔCt^ method.

### RNASeq

8 week old male C57BL/6J mice were injected with rmIL11 (200 mcg/kg) or an equivalent volume of saline (2 µL/kg). The left ventricle was excised and flash frozen 1, 3 or 6 hours after injection. Libraries were sequenced on a NextSeq 2000 to generate a minimum of 20 million paired end 60bp reads per sample.

Raw RNAseq data and gene-level counts have been uploaded onto the NCBI Gene Expression Omnibus database and will be made available with accession number (GSE240804).

### Single nuclei RNAseq

Single nuclei sequencing was performed on flash frozen LV tissue that was extracted from 8 week old male C57BL/6J mice 3 hours after injection with rmIL11 (200mcg/kg) or saline (2µL/kg). The tissue was processed according to standard protocols as previously described^20,21^. Nuclei were purified by fluorescent activated cell sorting and libraries were sequenced using HiSeq 4000 (Illumina, CA, USA) with a minimum depth of 20,000–30,000 read pairs per nucleus.

All single nuclei sequence data generated and analyzed in this study have been deposited as BAM files at the NCBI Gene Expression Omnibus database and will be made available upon request.

### ATAC Seq

8 week old male C57BL/6J mice were given an IP injection with rmIL11 (200mcg/kg) or saline. The heart was excised 3 hours after injection and flash-frozen tissue was sent to Active Motif to perform assay for transposase-accessible chromatin with sequencing (ATAC-seq) analysis.

### Protein Analysis

Protein extraction was performed on flash frozen tissue using ice-cold Pierce RIPA buffer (89901, ThermoFisher) supplemented with protease inhibitors (11697498001, Roche, Basel, Switzerland) and phosphatase inhibitors (4906845001, Roche). Tissue was lysed using a

Qiagen Tissue Lyser II with metallic beads for 3 mins at 30Hz. Protein quantification was performed using a Pierce bicinchoninic acid assay colorimetric protein assay kit (23225, ThermoFisher). 10-20µg of protein was loaded per well and run on a 4-12% bis-tris precast sodium-dodecyl sulfate page gel (NP0323BOX, Invitrogen). Semi-dry transfer was performed using the TransBlot Turbo transfer system (1704150, BioRad, CA, USA) and the membrane was blocked in 5% bovine serum albumin (A3803, Sigma-Aldrich, MO, USA). Primary antibodies raised against the following targets were used: signal transducer and activator of transcription 3 (STAT3) (4904S, Cell signalling technology (CST), MA, USA), pSTAT3 Tyr705 (9145L, CST), Extracelluar signal regulated kinase (ERK) (9101S, CST), pERK (4695S, CST), total c-Jun-N-terminal kinase (JNK) (sc-7345, Santa-Cruz, TX, USA), phospho-JNK (sc-6254, Santa-Cruz), green fluorescent protein (ab290, Abcam) Glyceraldehyde-3-phosphate dehydrogenase (GAPDH) (2118L, CST). Appropriate secondary horseradish peroxidase linked antibody was incubated for 1 hour with gentle agitation at room temperature and developed using chemiluminescence blotting substrate (1705061, BioRad or 34095, Thermofisher, depending on strength of signal).

### Cardiomyocyte extraction

CMs were extracted from the heart of 12 week old male C57BL/6J mice. Mice were deeply anaesthetized with ketamine and xylazine before the heart was harvested. Cells were incubated in Tyrode solution (1mM Ca, 1mM Mg) or Tyrode solution supplemented with rmIL11 (10 ng/mL) for 2 hours before recording. Cells were paced at 1Hz (10V, 10ms pulse width). Cell recordings were made using the Cytocypher high-throughput microscope (Cytocypher BV, Netherlands) and the automated cell finding system was used to identify and take recordings from 20 individual cells per heart per experimental condition. Calcium recordings were performed by incubating CMs with Fura 2AM dye (1uM) for 20 mins before fluorescent recordings were taken.

### Statistics

Statistical analyses were performed in GraphPad Prism V9.5.0 unless otherwise stated. Normality testing was performed using the Shapiro-Wilk test. Hypothesis testing for single comparisons was done using an unpaired two ways Student’s t-test for normally distributed data or by Mann-Whitney U test for non-normally distributed data.

Comparisons involving male and female mice were performed using a two-way analysis of variance (ANOVA) with Sidak’s multiple comparisons testing. Changes in expression over multiple time points were analysed using a one-way ANOVA with Sidak’s multiple comparisons testing. All graphs display the mean and standard error of the mean unless stated otherwise. P-values in RNA seq analysis were corrected for multiple testing using the false discovery rate (FDR) approach. A p-value and FDR of <0.05 was considered significant.

### Hierarchical testing of nested data

Statistical analysis of the data from high throughput microscopy of extracted cardiomyocyte experiments were analysed using a hierarchical statistical approach^22^. This approach tests for clustering within the data as may occur due to differences in the quality of myocyte preparation on different days. This uses a two-level random intercept model of linear regression. The analysis was performed using R-studio and the data was presented as the mean and standard deviation and effective n number taking the intraclass clustering into account.

Figures Graphs were prepared in GraphPad Prism V9.5.0 and R studio (Version 2023.03.0). Illustrations were created with Biorender.com and figures were arranged in Adobe Illustrator (Version 23.0.4.).

## Results

### Injection of rmIL11 to mice causes acute left ventricular dysfunction

Injection of rmIL11 to male C57BL/6J mice resulted in a tachycardic response (544±13 beats per minute (bpm), n=5) as compared to mice injected with saline (433±12 bpm, n=5) (Mann Whitney test, p=0.0079) **[Fig1A, B]**. Mice injected with rmIL11 injection had impaired LVEF (60.4%±3.1 vs 31.6±2.0, p<0.0001, n=5vs5), reduced GCS (−30.8%±2.3 vs −10.6±0.59, p<0.0001, n=5vs5) and reduced VTI in the aortic arch (36.8mm±1.9 vs 20.2±2.2, p<0.0004, n=5vs5) compared to mice injected with saline **[Fig1C-F]**. To serve as a related cytokine control an equivalent dose (200 mcg/kg) of rmIL6 was injected which had no detectable acute effects on cardiac function **[Fig1A-F & Suppl Fig S1A, B]**.

**Figure 1.**
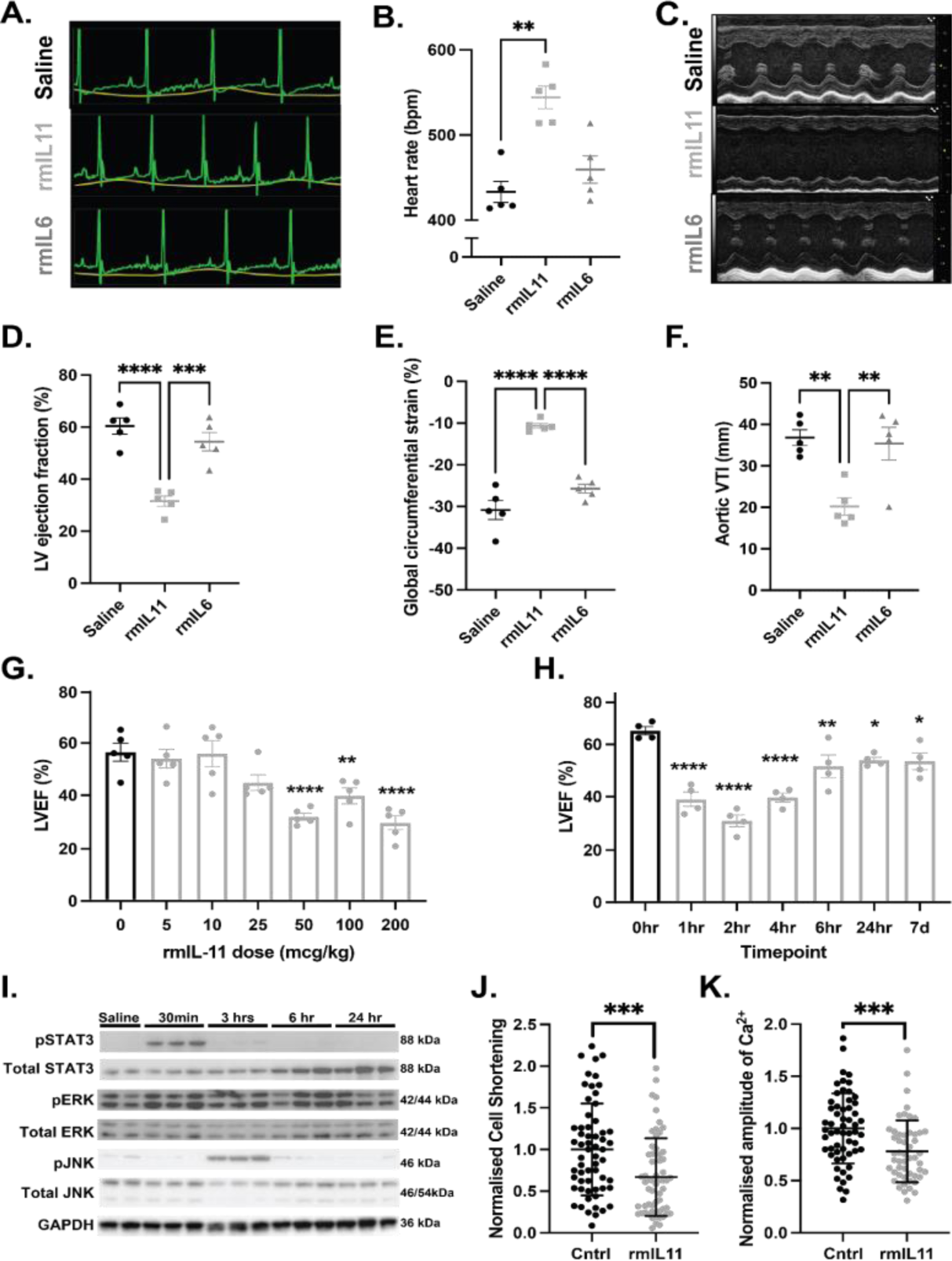
IL11 causes acute heart failure and impairs cardiomyocyte calcium handling. Male C57BL/6J mice were injected with rmIL11 (200 mcg/kg) (▪), rmIL6 (200 mcg/kg) (▴) or an equivalent volume of saline (2 µl/kg) (●). **(A)** Representative electrocardiogram traces were recorded under light anaesthesia, 2 hours after intraperitoneal (IP) injection of saline, rmIL11 or rmIL6. **(B)** Quantification of heart rate (n=5 per group). **(C)** Representative m-mode images from echocardiography performed 2 hours after injection of saline, rmIL11 or rmIL6. **(D)** Quantification of left ventricular ejection fraction (LVEF), **(E)** global circumferential strain and **(F)** velocity time integral at the aortic arch (VTI) in each group (n=5 per group). **(G)** LVEF 2 hours after IP injection of rmIL11 to male mice at 0, 5, 10, 25, 50, 100 and 200 mcg/kg (n=5 per dose). **(H)** LVEF at baseline, 1,2,4,6, and 24 hours, and 7 days after IP injection of rmIL11 (200 mcg/kg) (n=4 per timepoint). **(I)** Western blot of myocardial lysates from C57BL/6J male mice at baseline and 0.5, 3, 6 and 24 hours after IP rmIL11 injection (200 mcg/kg). Blots are probed for pSTAT3, total STAT3, pERK, total ERK, pJNK, total JNK and GAPDH. CMs isolated from male C57BL/6J mice were treated *in vitro* for 2 hours with media supplemented with rmIL11 (10 ng/mL) or non-supplemented media (Cntrl) (n=3 mice, 20 cells per mouse) and assessed for **(J)** contractility (effective n=9.7) and **(K)** the systolic change of intracellular calcium concentration (effective n=12). *Statistics: One-way ANOVA with Sidak’s multiple comparisons test. CM data: two level hierarchical clustering. Significance denoted as *p<0.05, **p<0.01, ***p<0.001, ****p<0.0001*.

Dosing studies revealed that the effects of rmIL11 on heart rate and left ventricular (LV) function were dose-dependent, consistent with physiological binding to and activation of the IlL11RA1 receptor. Cardiac impairment was evident at low doses and near-maximal effects were seen with a dose of 50 mcg/kg, which is the dose typically given daily to patients with thrombocytopenia post-chemotherapy **[Fig1G].** The effect of rmIL11 was rapid with a nadir in cardiac function at 2 hours post injection and recovery seen by 6 hours. However, recovery in cardiac function was incomplete and mild LV impairment persisted for up to 7 days following a single dose of rmIL11 **[Fig1H]**.

### IL11 causes impaired cardiomyocyte calcium handling

We next examined IL11 signalling pathways in cardiac extracts following rmIL11 injection, which revealed early and short-lived phosphorylation of signal transducer and activator of transcription 3 (p-STAT3) but no apparent ERK activation, which differs from acute signalling effects in the liver and other organs^23^ **[Fig1I & Suppl Fig S1C].** Phosphorylation of JNK is a stress related signalling pathway shown to be elevated in the mouse liver following IL11 treatment^23^. In the myocardium JNK was phosphorylated at the 3 hour time point post rmIL11 injection by which stage STAT3 phosphorylation was declining **[Fig1I & & Suppl Fig S1D]**.

The effect of IL11 directly on CMs was analysed *in vitro* by treating isolated adult mouse CMs with rmIL11 for 2 hours. CMs treated with rmIL11 demonstrated impaired contractility, as compared to control cells (Control: 1.00±0.177; rmIL11 (10 ng/mL): 0.669±0.150, p<0.000266) [**Fig1J]**. Intracellular calcium transients revealed blunting of the peak calcium concentration during systole in the presence of rmIL11 (Control: 1.00±0.0972; rmIL11: 0.781±0.0858, p<0.00019) **[Fig1K].**

### IL11 causes cardiac inflammation

The robust and early activation of STAT3 by IL11 led us to explore transcriptional changes which might occur acutely within the myocardium in response to IL11 injection. Bulk RNA sequencing was performed on LV tissue at 1, 3 and 6 hours following injection of rmIL11 and compared to controls injected with saline.

Extensive and significant transcription changes were apparent at all timepoints (**1hr,** Up:145, Down:27; **3hr,** Up:450, Down:303; **6hr:** Up: 268, Down:169; Log2FC+/-1, FDR<0.05). Genes differentially regulated included early upregulation of acute inflammatory genes (*Il6, Il1b* and *Il33*), chemotactic factors such as (*Ccl2* and *Cxcl1*) and CM stress markers *(Nppb, Cnn2, Ankrd1)* [**Fig2A, B**]. Kyoto Encyclopedia of Genes and Genomes (KEGG) analysis of the differentially expressed genes at the 1-hour time point revealed the TNFα, NF-κB and JAK/STAT signalling were among the most significantly enriched terms **[Fig2C & Suppl Fig S2A, C]**. A similar group of inflammatory terms were highlighted by Hallmark Gene Set Enrichment Analysis including TNFα signalling via NFκB, IL6 JAK/STAT and interferon-gamma signalling **[Fig2D & Suppl Fig S2B, D]**. These transcriptional changes show that IL11 drives an acute proinflammatory response in the heart that is associated with an impaired systolic function.

**Figure 2.**
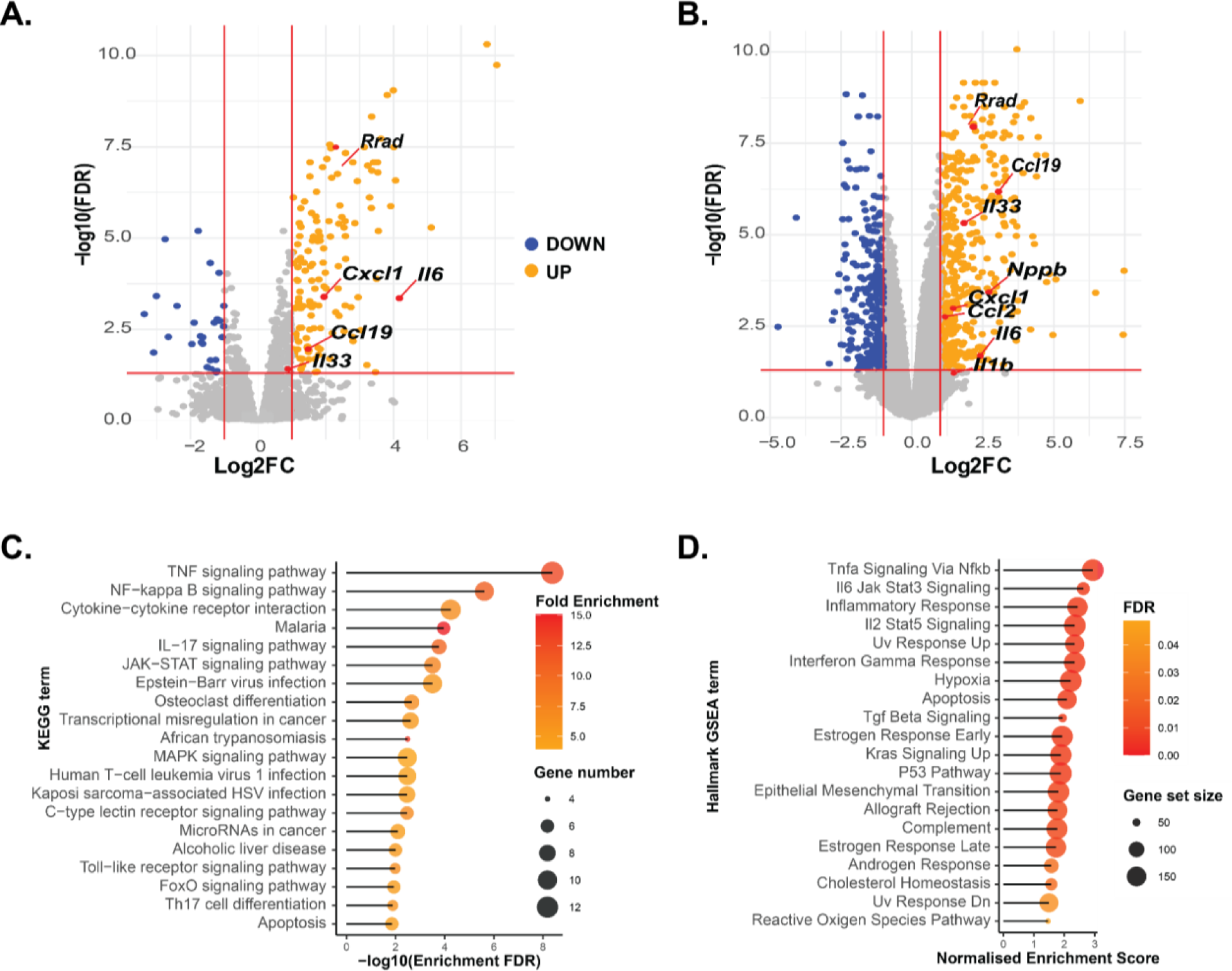
Transcriptional changes in the myocardium following rmIL11 injection. Volcano plot of all detected genes **(A)** 1 hour (n=3) and **(B)** 3 hours (n=4) after intraperitoneal injection of rmIL11 at 200 mcg/kg. Red lines are drawn at Log2Fc of 1 and −1 and FDR of 0.05. **(C)** Chart of most significantly enriched KEGG terms from at 1-hour post injection of rmIL11 ranked by FDR. **(D)** Gene set enrichment analysis of the most highly enriched Hallmark gene sets from RNAseq data at 1 hour after injection of rmIL11 ranked by normalised enrichment score.

On closer inspection, the L-type calcium channel inhibitor *Rrad* was identified as one of the most significantly upregulated genes at 1 hour (Fc:4.91, FDR:3.2e-8) and 3 hours (Fc:4.51, FDR:9.2e-9) post rmIL11 treatment **[Fig2A, B].** The *Rrad* protein product RAD-GTPase is a well-characterised and potent inhibitor of calcium current through L-type calcium channels^24,25^ and its acute upregulation may account for the changes in calcium transients seen in isolated CM preparations.

### Single nuclear sequencing reveals a cardiomyocyte stress signature

To examine cell-specific transcriptional responses and define any potential changes in cell populations, we performed single nucleus RNA-sequencing (snRNAseq) experiments on hearts 3 hours post rmIL11 injection **[Fig 3A, Suppl Fig S3A-C, S4A & Suppl Table 4]**. This revealed no significant change in cell populations overall, excluding immune cell infiltration at this early time point **[Fig 3B]**.

**Figure 3.**
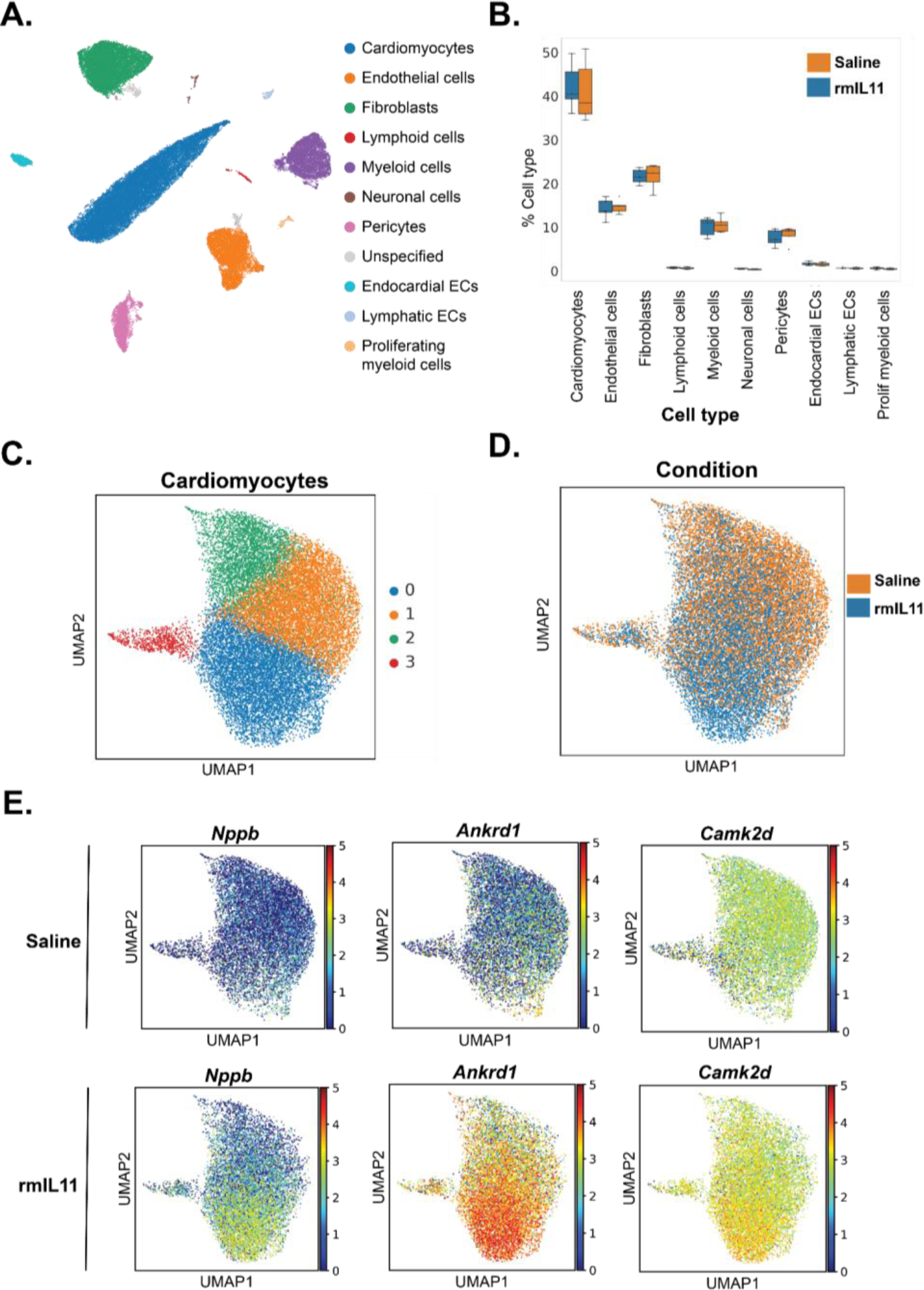
Single nuclear RNA sequencing reveals an IL11-induced cardiomyocyte stress signature. (A) UMAP embedding of all cell types from the left ventricle of male C57BL/6J mice 3 hours after intraperitoneal injection of rmIL11 (200 mcg/kg) or an equivalent volume of saline (n=5). **(B)** Comparison of cellular composition of the left ventricle in rmIL11 treated mice compared to saline treated mice. **(C)** UMAP embedding of cardiomyocyte fraction. 4 distinct clusters are identified based on gene expression. **(D)** UMAP embedding of cardiomyocytes annotated with the treatment group. **(E)** UMAP embedding of cardiomyocyte fraction of saline or rmIL11 treated cardiomyocytes annotated with relative expression of *Nppb, Ankrd1,* and *Camk2d*.

On closer analysis of CMs, this cell type segregated into four states with rmIL11-treated CM mostly localised to state 0 **[Fig 3C, D]**. This state is defined by the expression of cardiomyocyte stress factors including *Ankrd1*, *Ankrd23* and *Xirp2* **[Figure 3E & Suppl Fig S4B]***. Ankrd1* and *Ankrd23* are stress-inducible ankyrin repeat proteins which are elevated in dilated cardiomyopathies^26,27^. *Xirp2* encodes cardiomyopathy-associated protein 3 and is upregulated in CMs in response to stress^28,29^. Expression of *Nppb*, a canonical heart failure gene, was similarly elevated **[Fig 3E]**. Overall, the most enriched pathway from KEGG analysis of CM-specific differentially expressed genes, irrespective of state, was “Ribosome” with 93 out of 130 genes significantly upregulated (Fold enrichment:4.5, FDR:2.3e-46), perhaps related to the large effects of IL11 on protein translation and/or a pro-hypertrophic response of stressed CMs **[Suppl Fig S5]**^30^.

### ATAC-Seq highlights AP-1 family genes

To better understand the molecular changes induced by IL11 in the heart, we performed Assay for Transposase-Accessible Chromatin using sequencing (ATAC-seq) analysis. This methodology identifies regions of the genome undergoing epigenetic variation to make transcription factors binding sites more or less accessible.

Following IL11 administration, there were a large number of loci with variation in DNA accessibility (increased, 945; reduced, 445; shrunkenLog2FC:+/-0.3, Padj<0.1) **[Fig 4A & Suppl Table 5]**. The top twenty most differentially enriched regions **[Fig 4B, C]** include areas adjacent to *Camk2d, Ankrd1* and *Nppb,* stress and calcium handling genes that we had already found to be upregulated in CMs by snRNAseq [**Fig 3E**, **Fig 4B & Suppl Table 4**].

**Figure 4.**
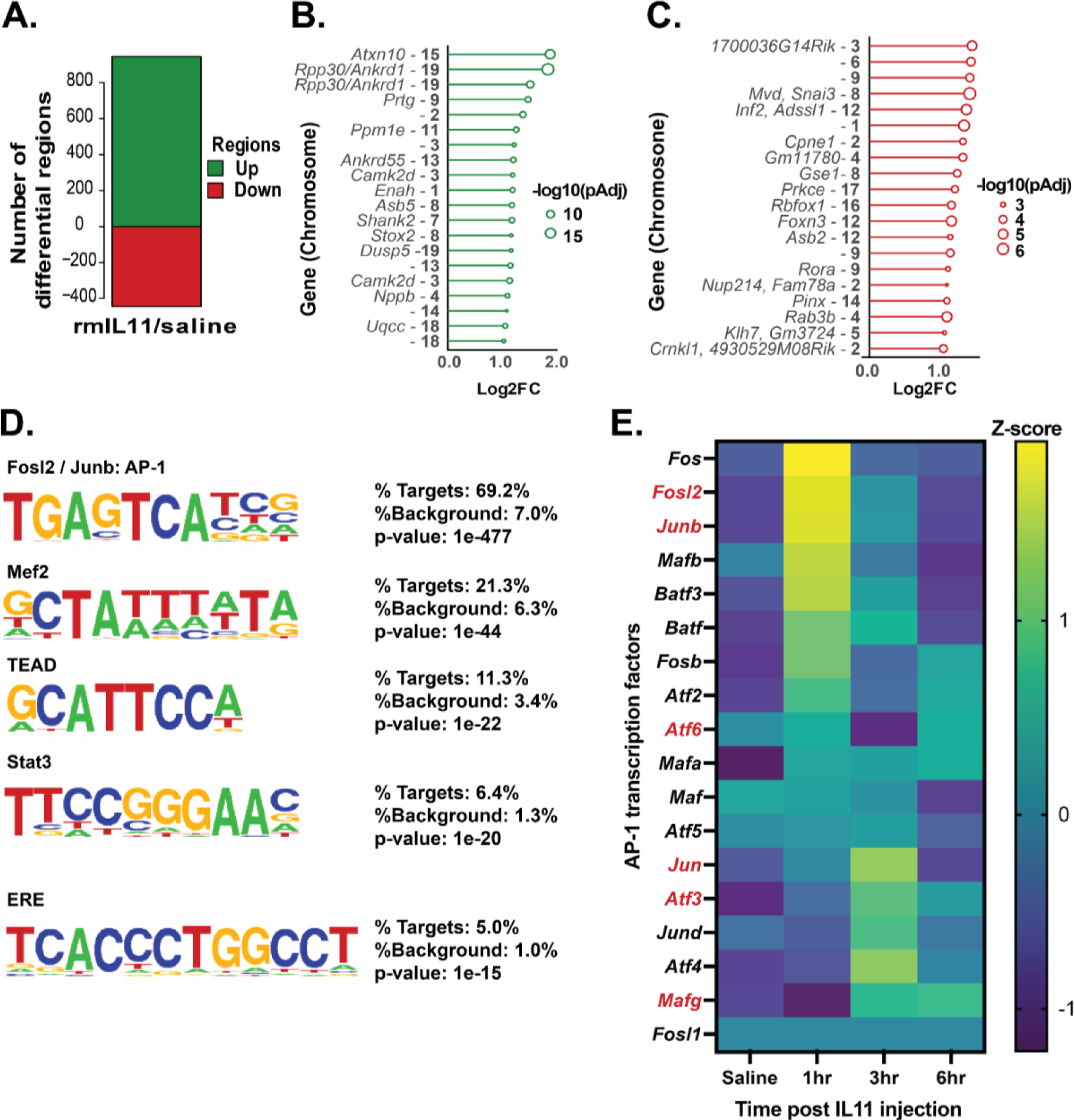
ATAC-Seq reveals a stress signature that occurs acutely in the myocardium after rmIL11 injection. **(A)** Number of positively and negatively enriched genomic regions identified by ATAC-Seq analysis of the myocardium 3 hours after injection of rmIL11 (n=4). **(B)** Top 20 most strongly enriched DNA regions in ATAC-seq analysis and adjacent genes, when present. **(C)** Top 20 most strongly negatively enriched DNA regions in ATAC-seq analysis and adjacent genes. **(D)** De novo Homer motif analysis of ATAC-seq data most highly enriched motifs in myocardial samples. **(E)** Heatmap of AP-1 transcription factor family members from bulk RNA sequencing data of myocardium at baseline, 1, 3 and 6 hours after rmIL11 injection. Genes differentially expressed in cardiomyocytes in single nuclear RNA sequencing data are highlighted in **red**.

DNA motif analysis of sequences captured by ATAC-seq, revealed the most enriched motifs after rmIL11 treatment were targets for FOSL2 and JUNB transcription factors **[Fig 4D & Suppl Table 6]**. These genes belong to the activator protein-1 (AP-1) transcription factor family, which is important for cardiomyocyte stress responses, cardiac inflammation and fibrosis.^31,32^ Notably, the STAT3 binding motif was also highly enriched.

We revisited our bulk RNA-seq data to examine the expression of the AP-1 transcription factor family transcripts after rmIL11 injection. This revealed that almost all of the AP-1 family transcripts are upregulated in the heart after rmIL11 **[Fig 4E]**. We then queried the snRNA-seq data and observed that *Fosl2, Junb, Atf6, Jun*, *Atf3* and *Mafg* are all significantly differentially expressed in cardiomyocytes following rmIL11 injection **[Fig 4E and Suppl Table 4]**.

### Viral-mediated CM-specific deletion of *Il11ra1*

To test whether the acutely negative inotropic effects of IL11 and the signature of increased CM stress are specifically mediated via IL11 activity in CMs, we proceeded to conditionally delete *Il11ra1* in CMs in the adult mouse. We used an adeno-associated virus serotype 9 (AAV9) vector to express *Tnnt2*-dependent *Cre*-recombinase in CMs of *Il11ra1* floxed mice, which effectively removed the floxed exons to generate mice with viral-mediated deletion of *Il11ra1* in CMs (vCMKO mice) **[Fig 5A, B]**. Effective transfection in the myocardium was confirmed by immunoblotting for GFP which is co-expressed with the *Cre*-recombinase [**Fig 5C**]. Notably, vCMKO mice had diminished myocardial p-STAT3 following injection of rmIL11, which demonstrates that IL11 activates JAK/STAT3 in CMs **[Fig 5C, D]**.

**Figure 5.**
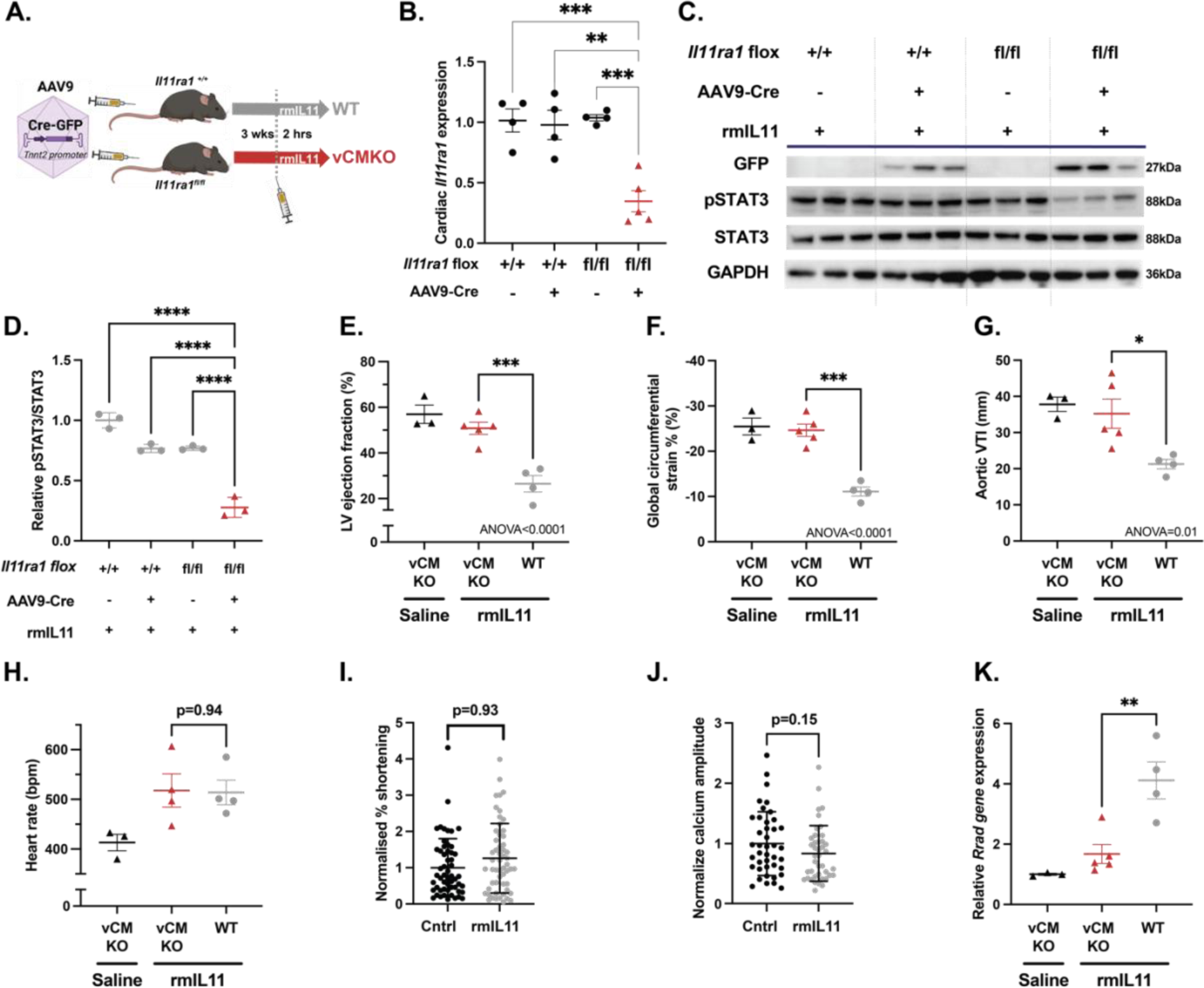
Viral-mediated *Il11ra1* deletion in adult cardiomyocytes protects against IL11-driven cardiac dysfunction. **(A)** Schematic of experimental design for AAV9 mediated delivery of *Tnnt2* promoter driven Cre-recombinase to male *Il11ra1^fl/fl^* or *Il11ra1^+/+^* mice. **(B)** QPCR of relative myocardial expression of *Il11ra1* in *Il11ra1^+/+^*or *Il11ra1^fl/fl^* injected with AAV9-Cre or vehicle. **(C)** Western blot from myocardial lysate following rmIL11 injection (200 mcg/kg) in *Il11ra1^+/+^* or *Il11ra1^fl/fl^*treated with either AAV9-Cre or saline (n=3). The membrane was probed with primary antibodies against GFP, pSTAT3, STAT3, and GAPDH. **(D)** Quantification of relative pSTAT3/STAT3 from (C). Echocardiographic assessment of vCMKO mice injected with rmIL11 (200 mcg/kg) (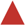) or saline (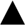) were compared to WT mice injected with rmIL11 (200 mcg/kg) (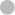). **(E)** Left ventricular ejection fraction, **(F)** global circumferential strain and **(G)** velocity time integral at the aortic arch and **(H)** heart rate were measured 2 hours after treatment (n=4). **(I)** Contractility (effective n = 53.6) and **(J)** peak calcium amplitude (effective n= 36.3) in CMs isolated from vCMKO mice and treated for 2 hours *in vitro* with rmIL11 containing media (10 ng/mL) or normal media (n=3 mice, 20 cells per mouse). **(K)** QPCR of myocardial expression of *Rrad* 2 hours vCMKO mice injected with rmIL11 (200 mcg/kg) (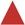) or saline (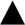) compared to WT mice injected with rmIL11 (200 mcg/kg) (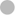) *Graphs CM data: mean ± standard deviation. Statistics: One-way ANOVA with Sidak’s multiple comparisons testing. CM data: two level hierarchical clustering. Significance denoted as *p<0.05, **p<0.01, ***p<0.001,****p<0.0001*.

As compared to mice injected with saline, WT mice injected with rmIL11 had impaired LVEF (WT+rmIL11: 26.5%±3.6), whereas vCMKO injected with rmIL11 had a mean LVEF (vCMKO+rmIL11: 50.8%±2.7) that was indistinguishable from saline-injected controls (vCMKO+saline: 57.0%±4.0) (n=3-5 per group) [**Fig 5E**]. Similar changes were seen in GCS (vCMKO+saline: −25.5%±1.9, vCMKO+rmIL11:-24.6%±1.4, WT+rmIL11: −11.1%±1.0, p<0.0001) and VTI in the aortic arch (vCMKO+saline: 37.8cm±1.9, vCMKO+rmIL11: 35.2cm±4.03, WT+rmIL11: 21.3cm±1.31, p<0.0371) **[Fig 5F, G]**. Interestingly, this experimental model still developed tachycardia following IL11 treatment, as seen in WT mice **[Fig 5H]**.

We performed experiments in CMs isolated from adult mice. Unlike CMs isolated from WT animals **[Fig 1J, K]**, CM from vCMKO mice did not have a reduction in cell shortening in response to stimulation with rmIL11, as compared to unstimulated cells (Cntrl: 1.0±0.11, vCMKO: 1.26±0.13, p=0.93, n=53.6). Similarly, peak calcium concentration was not blunted by rmIL11 in vCMKO CMs (Cntrl: 1.0±0.088, vCMKO: 0.83±0.076. p=0.15, n=36.3) **[Fig 5I, J].** As such, ILl1 effects in CMs are dependent on *Il11ra1* expression in CMs. Consistent with these changes in calcium signalling at a cellular level, qPCR of myocardial tissue of vCMKO mice prevented elevation of *Rrad* compared to control mice following rmIL11 injection **[Fig 5K]**.

### Germline deletion of *Il11ra1* in cardiomyocytes

We then used a complementary, germline deletion methodology to validate the CM-specific effects of IL11 seen in vCMKO mice by crossing *Il11ra1* flox mice with *Myh6*-Cre (m6CMKO) mice^33^ **[Fig 6A]**. This approach achieved a more pronounced and consistent knockdown of *Il11ra1* that enabled experiments to be scaled across sexes **[Fig 6B]**. As seen in the vCMKO strain, m6CMKO mice of both sexes had reduced p-STAT3 following rmIL11 injection, which further established effective *Il11ra1* locus recombination in this strain and reaffirms IL11-specific signalling in CMs **[Fig 6C, D]**.

**Figure 6.**
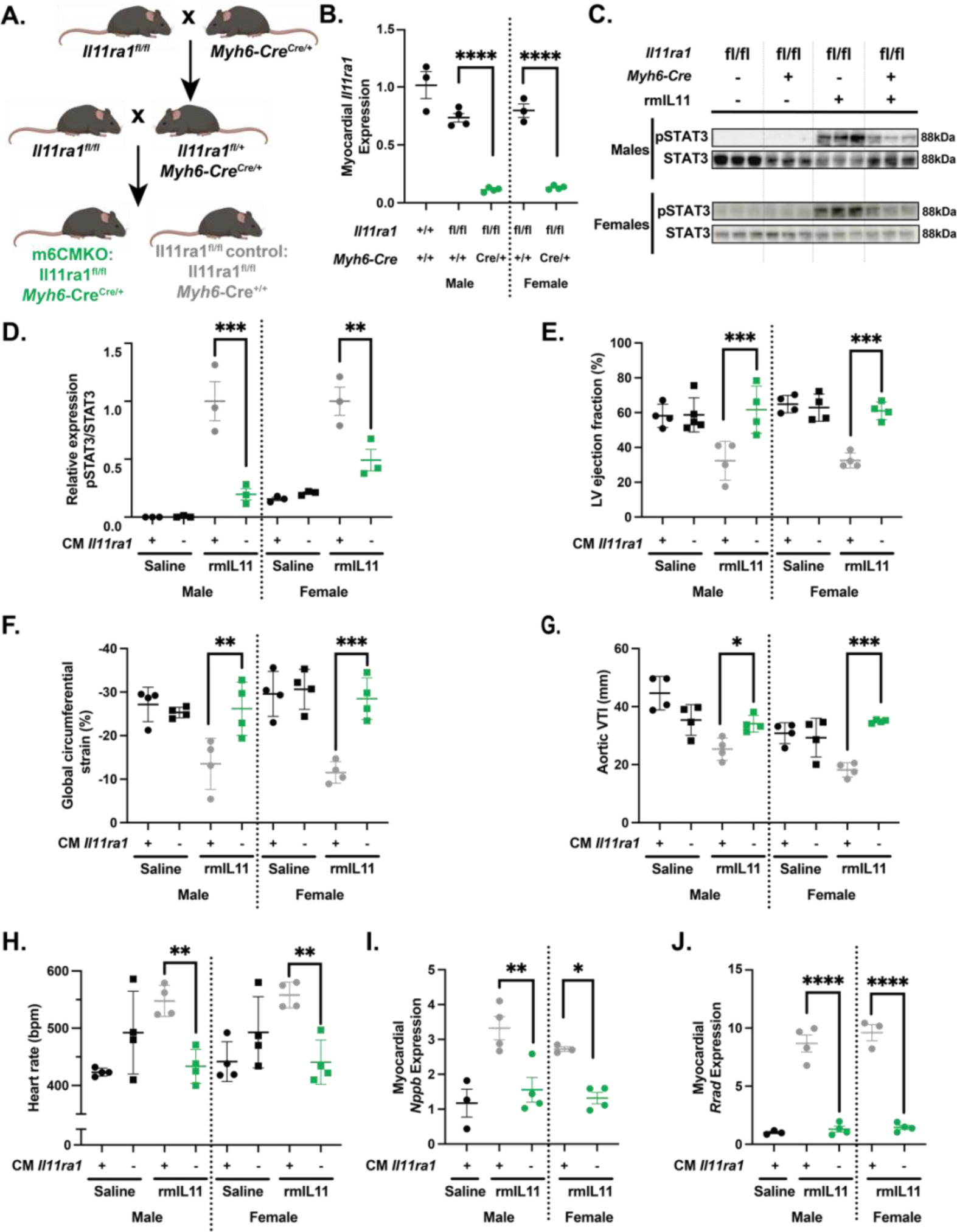
Germline deletion of *Il11ra1* in cardiomyocytes prevents IL11-induced cardiac toxicities. **(A)** Breeding strategy to generate m6CMKO mice and litter-mate *Il11ra1^fl/fl^*controls. **(B)** QPCR of *Il11ra1* gene expression in *Il11ra1^fl/fl^*controls and m6CMKO mice compared to male wild type C57BL/6J controls. (n=4) **(C)** Westerns blot of phospho-STAT3 and total STAT3 signalling in male and female *Il11ra1^fl/fl^* controls and m6CMKO mice with and without rmIL11 treatment. **(D)** Quantification of relative pSTAT and STAT3 expression. Male and female m6CMKO mice (CM *Il11ra -)* were treated with saline (▪) or rmIL11 (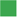) and compared to wild type mice (CM *Il11ra1* +) treated with saline (●) or rmIL11(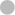) (n=4). **(E)** left ventricular ejection fraction, **(F)** global circumferential strain, **(G)** velocity time integral (VTI) in the aortic arch and **(H)** heart rate was measured 2 hours after rmIL11 injection. (n=4). QPCR analysis of relative expression of **(I)** *Nppb* and (**J)** *Rrad* in the myocardium following rmIL11 treatment of m6CMKO mice and *Il11ra1^fl/fl^* control mice (n=4). *Statistics: Comparison between groups by two-way ANOVA with Sidak’s multiple comparisons. p-values denoted as *<0.05, **<0.01, ***<0.001, ****<0.0001)*.

Having established the m6CMKO strain, we examined the effects of rmIL11 on cardiac function in these mice **[Suppl Table 7]**. When injected with rmIL11, *Il11ra1^fl/fl^* control mice had significantly reduced LVEFs whereas the LVEF of m6CMKO was similar to that of m6CMKO mice injected with saline **[Fig 6E]**. Similarly, following rmIL11 injection, GCS and VTI in the aortic arch were impaired in control mice expressing *Il11ra1* but not in m6CMKO mice **[Fig 6F, G].** It was evident that the molecular and cardiovascular phenotypes of m6CMKO mice injected with rmIL11 largely replicated those observed in the vCMKO mice. However, different to the vCMKO strain, the m6CMKO mice were protected against IL11-induced tachycardia. **[Fig 6H]**.

In molecular studies of myocardial extracts, *Nppb* was upregulated in *Il11ra^fl/fl^* control mice in response to rmIL11 injection, but this was not seen in m6CMKO mice **[Fig 6I]**. Similarly, following rmIL11 injection the L-type calcium channel inhibitor *Rrad* was upregulated in *Il11ra^fl/fl^* controls but not in m6CMKO mice **[Fig 6J].**

### JAK inhibition protects against IL11-induced cardiac dysfunction

Canonical IL11 signalling through the IL11RA/gp130/JAK/STAT3 pathway has recently been implicated in the acute pro-inflammatory effects of IL11^34^ and activation of STAT3 in the heart was immediate and pronounced following IL11 injection [**Fig 1I**]. To determine the functional relevance of JAK/STAT3 activation in the heart we pretreated mice with ruxolitinib (30 mg/kg), which inhibits JAK1/2 activation, prior to injection of rmIL11 **[Fig 7A]**.

**Figure 7.**
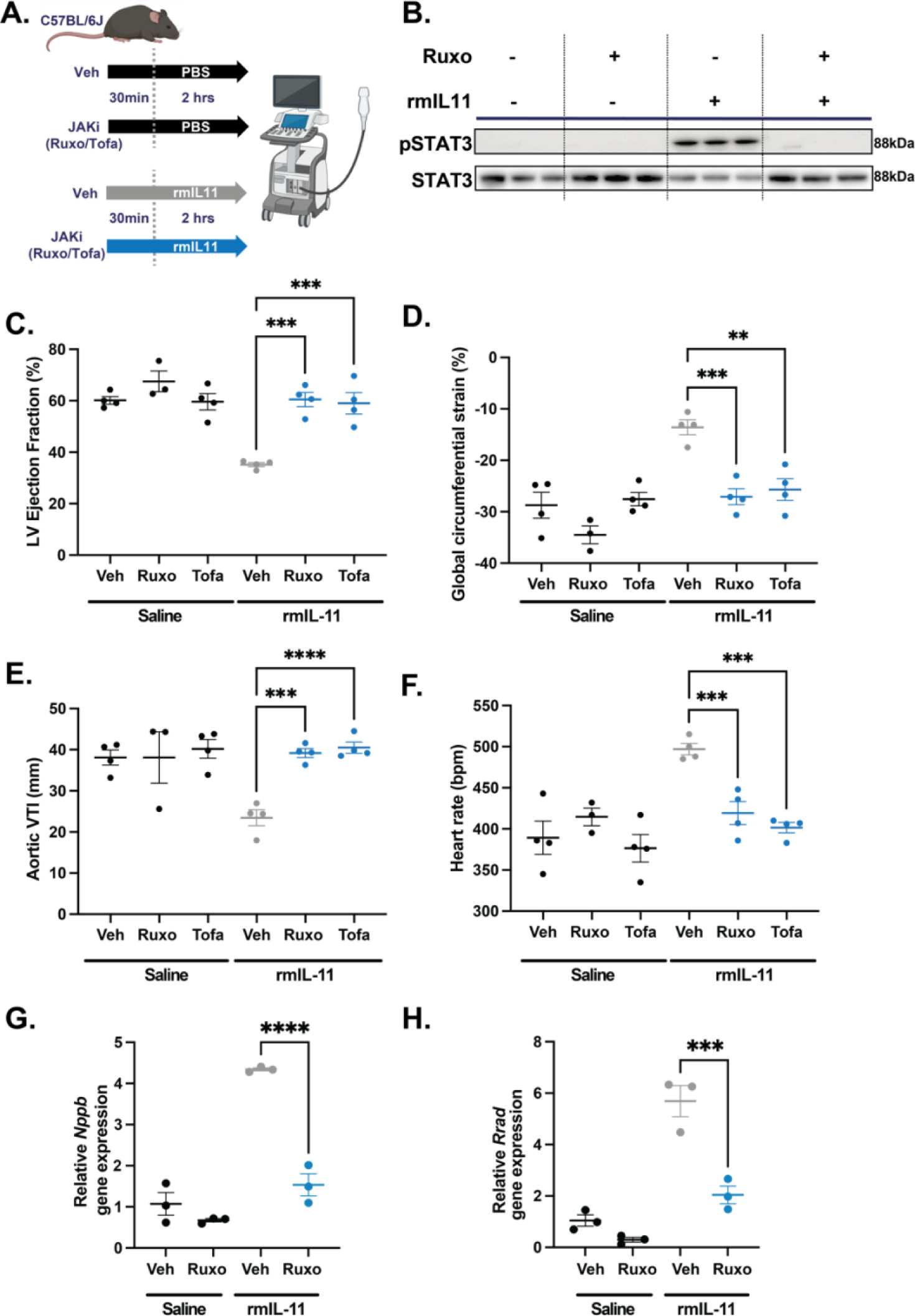
The acute toxic effects of rmIL11 are mediated via JAK/STAT signalling. **(A)** Schematic of the pretreatment of wild type male C57BL/6J mice with JAK inhibitor (JAKi) or vehicle (Veh) 30 mins before administration of recombinant mouse IL11 (rmIL11) or saline. **(B)** Western blot of myocardium lysate in mice treated with a combination of ruxolitinib (30 mg/kg) or vehicle and saline or rmIL11. Membranes have been probed for pSTAT3 and STAT3 (n=3). Mice treated with a combination of vehicle (Veh), ruxolitinib (Ruxo) or tofacitinib (Tofa) and either saline or rmIL11 had an echocardiogram performed 2 hours after treatment which measured **(C)** left ventricular ejection fraction, **(D)** global circumferential strain, **(E)** velocity time integral (VTI) in the aortic arch and **(F)** heart rate (n=4). QPCR of myocardial tissue from combinations of Ruxo and rmIL11 treatments of **(G)** *Nppb* expression and **(H)** *Rrad* expression (n=3). *Statistics: Comparison between groups by one-way ANOVA with Sidak’s multiple comparisons test. Significance denoted as denoted *p<0.05, **p<0.01,****p<0.0001*.

We confirmed that administration of ruxolitinib at 30 mg/kg prevented activation of JAK/STAT3 signalling by immunoblotting **[Fig 7B]**. Having established the efficacy of ruxolitinib we studied its effect on cardiac physiology in 8 week old wild type male mice injected with rmIL11. Ruxolitinib alone had no effect on LV function **[Fig 7C]**. Following injection of rmIL11, and as compared to buffer injected controls, mice pretreated with ruxolitinib had better LVEF (60.5%±2.79 vs 35.2%±0.79; p=0.0005), GCS (−27.1%±1.56 vs - 13.6%±1.44 vs, p=0.0009) and aortic VTI (39.2cm±10.9 vs 23.4cm±1.92, p=0.0001) **[Fig 7C-E]**). Ruxolitinib pretreatment also prevented rmIL11-induced tachycardia (497±6.8 vs 419±14.1, p=0.0008) **[Fig 7F]**. As seen with m6CMKO, JAK inhibition prevented stress associated transcriptional changes in the heart of *Nppb* and *Rrad* **[Fig 7G, H]**.

To exclude off-target effects and to replicate findings, the study was repeated with a second JAK inhibitor (tofacitinib, 20 mg/kg). As seen with ruxolitinib, pretreatment with tofacitinib protected against the varied deleterious effects of IL11 on cardiac function compared to vehicle treated controls: LVEF (59.0±4.16, p=0.0007), GCS (−25.7±2.10, p=0.002), VTI in the aortic arch (40.5±1.36, p<0.0001), and tachycardia (401±6.23, p=0.0002). **[Fig 7C-E]**.

## Discussion

In some healthcare systems, rhIL11 is used routinely to increase platelet counts in patients with thrombocytopenia but this can cause serious cardiac complications that are unexplained and thought non-specific. RhIL11 has also been trialled in a different context, as a cytoprotective agent, in patients across a range of other medical conditions (e.g. colitis, myocardial infarction, arthritis, cirrhosis), **[Table 1 & Supl Table 1]** as IL11 was previously thought to be anti-inflammatory and anti-fibrotic^16^. As such, many thousands of patients have received, and continue to receive, rhIL11 in clinical trials and as part of routine medical care. Long-acting formulations of rhIL11 have recently been devised and new clinical trials of rhIL11 are proposed^5^.

The cardiac side effects of rhIL11, while unexplained, have long been recognised and a small clinical trial was initiated in 2009 to determine if rhIL11 (50 mcg/kg) had an effect on cardiac conduction (NCT00886743). This trial was terminated prematurely at the request of the sponsor and no formal conclusions were made. Other studies looking at the effects of injection of human IL11 to adult rats showed no effects on cardiac phenotypes and studies of human atrial myocytes were similarly negative^35,36^. We suggest that, for these reasons, the severe cardiac side effects of rhIL11 therapy have been explained away as indirect, non-specific and thus sidelined^36^.

We found that injection of species-matched rmIL11 to mice caused acute and dose-dependent LV impairment that was mediated via IL11’s action in IL11RA1 expressing CMs. CMs exposed to IL11 displayed perturbed calcium handling, upregulated cellular stress factors (*Ankrd1, Ankrd23, Xirp2, Nppb)* and had exhibited increased inflammation (TNFα, NFκB and JAK/STAT). These findings further redress the earlier literature on IL11 activity in the heart where it was believed to be anti-fibrotic^14^, which appears inaccurate^30^, and that it was cytoprotective in CMs^13–15^, which we challenge here. In retrospect, the use of rhIL11 in a clinical trial of patients with myocardial infarction was likely ill-founded^6^.

The powerful enrichment of the AP-1 family of transcription factors following rmIL11 injection, seen in bulk RNA-seq, snRNA-seq and ATAC-seq was unexpected and likely has detrimental effects in the mouse heart^31,37^. AP-1 family activation is not immediately downstream of IL11:IL11RA:gp130 signalling and thus, the early IL11-stimulated activation of JAK/STAT3 likely primes CMs to upregulate, activate and respond to AP-1 transcription factors. In the injured zebrafish heart, AP-1 contributes to sarcomere disassembly and regeneration^38^, which is IL11-dependent^39^, providing an evolutionary context for IL11-mediated effects in the heart^40^.

Our use of two mouse models of CM-specific *Il11ra1* deletion show and replicate that the effects of rmIL11 on cardiac function are via direct cardiotoxic effects on CMs and are not explained by changes in circulating volume, as has previously been suggested^36^. The models used in this study involved the administration of a single dose of rmIL11 however in clinical practice, courses of therapy can involve daily infusions of rhIL11 for up to 21 days between chemotherapy cycles which are likely to compound the effect on the heart, specifically on fibrotic pathologies that are slow to establish^30^.

There are several limitations to our study. The discrepancy between the tachycardia seen in vCMKO but not m6CMKO mice suggests a direct effect of IL11 on sinoatrial node, which is differentially deleted for *Il11ra1* between the models, was not explored. We did not determine the putative roles of *Rrad* or *Camk2d* in IL11-induced contractile dysfunction, which should be investigated more fully in follow-on studies. Effects on human CMs were not examined, although we have observed conserved effects of IL11 on multiple cellular phenotypes in varied human and mouse cell types (e.g. fibroblasts, hepatocytes, epithelial cells)^16,23^. Whether endogenous IL11 is toxic to CMs and negatively inotropic in heart failure syndromes is not known and we cannot extrapolate from the data seen with acute, high dose injection of recombinant protein. The cardiac side effects associated with IL11 include arrhythmias (notably atrial fibrillation and flutter) that we did not study here.

In conclusion, we show for the first time that IL11 injection causes IL11RA-dependent, CM-specific toxicities and acute heart failure. These data may explain the serious cardiac side effects that occur with rhIL11 therapy, which have been overlooked. Our findings question the ongoing use of rhIL11, and its further development^5^, in patients with thrombocytopenia while identifying novel toxic effects of IL11 in the cardiomyocyte compartment of the heart.

## Supporting information

Uncropped blots

## Abbreviations

AAV9: Adeno-associated virus serotype 9
ANOVA: Analysis of variance
AP-1: Activator protein 1
ATACseq: Assay for transposase-accessible chromatin with sequencing
Bpm: Beats per minute
CM: Cardiomyocyte
ECG: Electrocardiogram
EGFP: Enhanced green florescent protein
ERK: Extracellular signal regulated kinase
FDR: False discovery rate
FOSL2: FOS like 2
GAPDH: Glyceraldehyde-3-phosphate dehydrogenase
GCS: Global circumferential strain
GSEA: Gene set enrichment analysis
HR: heart rate
IL6: Interleukin 6
IL11: Interleukin 11
IL11RA1: Interleukin 11 receptor A1
IP: Intraperitoneal
JAK: Janus kinase
JNK: c-Jun N-terminal kinase
KEGG: Kyoto encyclopaedia of genes and genomes
LV: Left ventricle
LVEF: Left ventricular ejection fraction
PBS: Phosphate buffered saline
PCR: Polymerase chain reaction
PSAX: Parasternal short axis
QPCR: Quantitative polymerase chain reaction
rhIL11: Recombinant human interleukin 11
RIPA: Radioimmunoprecipitation assay buffer
rmIL6: Recombinant mouse interleukin 6
rmIL11: Recombinant mouse interleukin 11
SEM: Standard error of the mean
STAT: Signal transducer and activator of transcription
TNFα: Tumour necrosis factor-alpha
UMAP: Uniform Manifold Approximation and Projection
vCMKO: Viral mediated cardiomyocyte Il11ra1 knockout
VTI: Velocity time integral
WT: Wild type

