## Supplemental Materials for "Interleukin 11 therapy causes acute heart failure and its use in patients should be reconsidered"

### Supplementary Materials

#### Detailed Methods

**RNAseq**

RNA quality for RNAseq was assessed using the Agilent 2100 RNA 6000 Nano assay and RNASeq libraries prepared using the NEBNext Ultra II Directional RNA Library Prep Kit with NEBNext Poly(A) mRNA Magnetic Isolation Module following manufacturer's instructions. Library quality was evaluated using the Agilent 2100 High-Sensitivity DNA assay, and concentrations measured using the Qubit™ dsDNA HS Assay Kit.

Raw reads were aligned to the mouse reference genome GRCm39 (Ensembl release 106) using the STAR[41] software v2.7.10a with default parameters. Aligned reads were assigned to genes to give a gene-level count matrix using the featureCounts tool[42] as part of the Rsubread package v2.10.4. Differential gene expression analysis was performed using the edgeR[43] software v3.38.1 following the author's recommended analysis pipeline utilising a quasi-likelihood negative binomial generalized log-linear model.

One of the 1 hour samples was excluded after randomisation as no phosphorylation of STAT3 was seen on western blot analysis and there were no significantly differentially expressed genes compared to saline injected mice on RNA seq analysis. It was assumed that the rmIL11 intraperitoneal injection was ineffective and may have been intraluminal. Therefore, this sample was excluded from further analysis and the 1 hour RNAseq group has 3 rather than 4 biological replicates.

Gene set enrichment analysis (GSEA) was carried out on fold change ranked gene lists from male mice 1, 3, and 6 hours after injection of rmIL11 and compared to injection with saline. The data was imputed into the gene set enrichment analysis software (version 4.3.2, Broad Institute, San Diego, USA & UC San Diego, USA) with MSigDB Hallmark gene sets.[44,45]

KEGG pathway analysis was performed using differentially expressed gene lists ($Log_2FC$+/-1, FDR<0.05)[46,47]. Gene lists were imputed into the ShinyGO software (v0.77, South Dakota State University, USA)[48] and the top 20 KEGG pathways ranked by enrichment false discovery rate (FDR) values were visualised.

**Single nuclei RNAseq**

Single nuclei were obtained from flash-frozen tissues using mechanical homogenization in homogenization buffer (250 mM sucrose, 25 mM KCl, 5 mM MgCl2, 10mM Tris-HCl, 1 mM dithiothreitol (DTT), 1× protease inhibitor, 0.4 U μl−1 RNaseIn, 0.2 U μl−1 SUPERaseIn, 0.1% Triton X-100 in nuclease-free water) (Litvinukova et al., 2018). Nuclei were stained with NucBlue Live ReadyProbes Reagents (ThermoFisher) and Hoechst-positive single nuclei were purified by fluorescent activated cell sorting using influx, XDP or FACSAria (BD Biosciences, Heidleberg, Germany).

Nuclei purification and integrity was verified under a microscope, and nuclei were further processed using the Chromium Controller (10X Genomics, CA, USA) according to the manufacturer's protocol. Single nuclei were manually counted by Trypan blue exclusion. Nuclei suspension was adjusted to 400–1,000 cells per microlitre and loaded on the Chromium Controller (10X Genomics) with a targeted nuclei recovery of 4,000–10,000 per reaction. 3′

gene expression libraries were prepared according to the manufacturer's instructions Chromium Single Cell Reagent Kits (10X Genomics). Libraries were sequenced using HiSeq 4000 (Illumina, CA, USA) with a minimum depth of 20,000–30,000 read pairs per nucleus.

#### ATACSeq

The tissue was manually disassociated, isolated nuclei were quantified using a haemocytometer, and 100,000 nuclei were tagmented as previously described,[49] with some modifications based on Corces et al[50] using the enzyme and buffer provided in the Nextera Library Prep Kit (Illumina, CA, USA). Tagmented DNA was then purified using the MinElute PCR purification kit (Qiagen), amplified with 10 cycles of PCR, and purified using Agencourt AMPure SPRI beads (Beckman Coulter, CA, USA). The resulting material was quantified using the KAPA Library Quantification Kit for Illumina platforms (KAPA Biosystems, MA, USA), and sequenced with PE42 sequencing on the NextSeq 500 sequencer (Illumina).

For analysis, reads were aligned using the BWA algorithm (mem mode; default settings). Duplicate reads were removed, only reads mapping as matched pairs and only uniquely mapped reads (mapping quality >= 1) were used for further analysis. Alignments were extended in silico at their 3'-ends to a length of 200 bp and assigned to 32-nt bins along the genome. The resulting histograms (genomic "signal maps") were stored in bigWig files. Peaks were identified using the MACS 3.0.0 algorithm at a cut-off of p-value 1e-7, without control file, and with the nomodel option. Peaks that were on the ENCODE blacklist of known false ChIP-Seq peaks were removed. Signal maps and peak locations were used as input data to Active Motifs proprietary analysis program, which creates Excel tables containing detailed information on sample comparison, peak metrics, peak locations and gene annotations. For differential analysis, reads were counted in all merged peak regions (using Subread), and the replicates for each condition were compared using DESeq2.

**Cardiomyocyte Isolation**

The chest was opened and the heart and aorta were transferred to ice-cold perfusion buffer (113mM NaCl, 4.7mM KCl, 0.6mM $KH_2PO_4$, 0.6mM $Na_2HPO_4$, 1.2mM $MgSO_4$-7H2O, 12mM $NaHCO_3$, 10mM $KHCO_3$, HEPES Na Salt 0.922mM, Taurine 30mM, butanedione monoxime 10mM, glucose 5.5mM, pH 7.4). Excess material was dissected away from the heart and the ascending aorta cannulated and placed on retrograde perfusion Langendorff apparatus. The heart was perfused with warmed perfusion buffer for 4 mins and then digestion buffer (0.2mg/mL Liberase ™, (05401127001, Sigma Aldrich), 5.5mM Trypsin (15090046, Invitrogen), 0.005U/mL DNase (79256, Qiagen), 12.5uM $CaCl_2$) until fully digested. Subsequently, the heart was minced and gently pipetted until a uniform suspension was formed. Cells were filtered through sterile gauze and resuspended in perfusion buffer. Sequential calcium reintroduction was performed in 3 steps up to 1mM (62uM, 212uM and 1mM). CMs were plated on laminin and allowed to equilibrate for 1 hour in M199 media supplemented with 2mM L-carnitine, 5mM creatine, and 5mM taurine before stimulation with rmIL11.

### Supplementary Tables

| NCT Number / PMID | Title | Study Completion / Publication Date | Number of Patients | Status |
|---|---|---|---|---|
| **Chemotherapy Induced Thrombocytopenia** | | | | |
| NCT03823079 | Comparison of Interleukin-11 and rhTPO for Recurrent Colorectal Cancer Patients With Thrombocytopenia | Feb-20 | 50 | Unknown status |
| NCT01663441 | A Phase IIIa Study of Genetically Modified Recombinant Human Interleukin-11 | Aug-17 | 62 | Completed |
| NCT02314273 | Effect of rhIL-11 in Patients With Thrombocytopenia for Childhood ALL | Apr-14 | 120 | Completed |
| NCT00886743 | Study Evaluating The Effects Of Oprelvekin On Cardiac Repolarization In Subjects With Chemotherapy Induced Thrombocytopenia | Dec-15 | 19 | Terminated |
| NCT00493181 | Interleukin 11, Thrombocytopenia, Imatinib in Chronic Myelogenous Leukemia (CML) Patients | Jan-11 | 8 | Completed |
| PMID: 12839693 | Recombinant human interleukin-11 in the prevention of chemotherapy-induced thrombocytopenia | May-03 | 118 | Completed |
| PMID: 12717398 | Recombinant human interleukin-11 improves thrombocytopenia in patients with cirrhosis | Mar-01 | 10 | Completed |
| PMID: 22041866 | Multicenter, randomized study of genetically modified recombinant human interleukin-11 to prevent chemotherapy- Nov-11 induced thrombocytopenia in cancer patients receiving chemotherapy | Aug-12 | 88 | Completed |
| PMID: 21122391 | Therapeutic effect of interleukin-11 on thrombocytopenia in patients with hematologic malignancies after chemotherapy | Sep-10 | 70 | Completed |
| PMID: 19236785 | Effect of recombinant human interleukin 11 on recovery of platelets after peripheral blood hematopoietic stem cell transplantation | Feb-09 | 20 | Completed |
| PMID: 17611320 | Effect of recombinant human interleukin 11 on the platelet after hematopoietic stem cell transplantation in patients with leukemia | Jun-07 | 24 | Completed |

| | | | | |
|---|---|---|---|---|
| PMID: 17261503 | A multicenter randomized, double-blind, placebo-controlled late-phase II/III study of recombinant human interleukin 11 in acute myelogenous leukemia | Jan-07 | 101 | Completed |
| PMID: 17147124 | Effectiveness and safety of recombinant human interleukin-11 in the treatment of chemotherapy-induced thrombocytopenia | Jul-06 | 32 | Completed |
| PMID: 16413056 | Phase II trial of subcutaneous recombinant human interleukin 11 with subcutaneous recombinant human granulocyte-macrophage colony stimulating factor in patients with acute myeloid leukemia (AML) receiving high- dose cytarabine during induction: ECOG 3997 | Jul-06 | 34 | Completed |
| PMID: 16117903 | Phase II clinical trial on China manufactured recombinant human interleukin-11 derivative in the prevention and treatment of chemotherapy-induced thrombocytopenia | Jun-05 | 100 | Completed |
| PMID: 15853214 | A clinical study of YM 294 (rhlL-11) in patients with gynecologic cancer | Apr-05 | - | |
| PMID: 15853215 | An early phase II clinical study of YM 294 (rhlL-11) in patients with solid tumors and malignant lymphoma | Apr-05 | - | Completed |
| PMID: 15606549 | Phase I/II dose escalation study of recombinant human interleukin-11 following ifosfamide, carboplatin and etoposide in children, adolescents and young adults with solid tumours or lymphoma: a clinical, haematological and biological study | Jan-05 | 47 | Completed |
| PMID: 12478901 | Clinical study of rhIL-11 for prevention and treatment of chemotherapy-induced thrombocytopenia | Aug-02 | 109 | Completed |
| PMID: 9923411 | A randomized trial of recombinant human interleukin-11 following autologous bone marrow transplantation with peripheral blood progenitor cell support in patients with breast cancer | Nov-98 | 80 | Completed |
| PMID: 9363868 | Randomized placebo-controlled study of recombinant human interleukin-11 to prevent chemotherapy-induced thrombocytopenia in patients with breast cancer receiving dose-intensive cyclophosphamide and doxorubicin | Nov-97 | 77 | Completed |
| **Immune Thrombocytopenic Purpura** | | | | |
| PMID: 26524046 | Therapeutic Efficacy of Multigly-Cosidorum Tripterygium Combined with rhIL-11 for Immune Thrombocytopenia | Oct-15 | 75 | Completed |
| PMID: 18401155 | Interleukin-11 for Treatment of Hepatitis C-Associated ITP | Jan-08 | 12 | Completed |
| PMID: 18928632 | Curative effect of rhIL-11 on 26 patients with chromic idiopathic thrombocytopenic purpura | Oct-08 | 26 | Completed |
| PMID: 11279623 | A pilot study of rhuIL-11 treatment of refractory ITP | Mar-01 | 7 | Completed |

| | | | | |
|---|---|---|---|---|
| **Coagulopathy** | | | | |
| NCT00994929 | Phase II Prospective Open-Label Trial of Recombinant Interleukin-11 in DDAVP-Unresponsive Von Willebrand Disease and Mild or Moderate Hemophilia A | Apr-12 | 9 | Completed |
| NCT00524225 | IL-11 in Adults With Von Willebrand Disease Undergoing Surgery | Jun-12 | 3 | Terminated |
| PMID: 21833452 | IL-11 in Women With Von Willebrand Disease and Refractory Menorrhagia | Oct-11 | 7 | Completed |
| NCT00151125 | Phase II Study of IL-11 (Neumega) in Von Willebrand Disease | Dec-07 | 12 | Completed |
| **Inflammatory Bowel Disease** | | | | |
| PMID: 16635225 | Randomized, double blind controlled trial of subcutaneous recombinant human interleukin-11 versus prednisolone in active Crohn's disease | Apr-06 | 51 | |
| NCT00038922 | Study Evaluating rhIL-11 in Left-Sided Ulcerative Colitis | Jan-00 | | Terminated |
| NCT00040521 | Study Evaluating rhIL-11 in Active Crohn's Disease | Oct-03 | | Completed |
| PMID: 10381910 | Preliminary evaluation of safety and activity of recombinant human interleukin 11 in patients with active Crohn's disease | Jul-99 | 76 | Completed |
| PMID: 11876692 | Randomized, controlled trial of recombinant human interleukin-11 in patients with active Crohn's disease | Mar-98 | 148 | Completed |
| **Malignancy** | | | | |
| PMID: 29798418 | Effect of recombinant human interleukin-11 treatment on prognosis of patients with radiochemoradiotherapy in the treatment of nasopharyngeal carcinoma | Jun-17 | 78 | Completed |
| PMID: 21418991 | Effects of recombinant human interleukin 11 on hematological malignancy after allogeneic hematopoietic cell transplantation | Jun-10 | 48 | Completed |
| NCT00012298 | Radiolabeled Monoclonal Antibody Plus Rituximab With and Without Filgrastim and Interleukin-11 in Treating Patients With Relapsed or Refractory Non-Hodgkin's Lymphoma | Apr-10 | 81 | Terminated |

| | | | | |
|---|---|---|---|---|
| PMID: 16644288 | Modulation of the systemic inflammatory response by recombinant human interleukin-11: A prospective randomized placebo controlled clinical study in patients with hematological malignancy | Aug-06 | 40 | Completed |
| PMID: 12036860 | Gemtuzumab ozogamicin with or without interleukin 11 in patients 65 years of age or older with untreated acute myeloid leukemia and high-risk myelodysplastic syndrome: comparison with idarubicin plus continuous-infusion, high-dose cytosine arabinoside | Jun-02 | 51 | Completed |
| PMID: 12559860 | Recombinant human interleukin 11 and bacterial infection in patients with [correction of] haematological malignant May-02 disease undergoing chemotherapy: a double-blind placebo-controlled randomised trial | Jan-03 | 40 | Completed |
| **Bone Marrow Failure** | | | | |
| PMID: 17071475 | Phase II study of low-dose interleukin-11 in patients with myelodysplastic syndrome | Apr-06 | 35 | Completed |
| PMID: 11689585 | Pilot study of low-dose interleukin-11 in patients with bone marrow failure | Nov-01 | 16 | Completed |
| PMID: 11464987 | Feasibility study of IL-11 and granulocyte colony-stimulating factor after myelosuppressive chemotherapy to mobilize peripheral blood stem cells from heavily pretreated patients | Dec-98 | 42 | Completed |
| **Psoriasis** | | | | |
| PMID: 10587516 | Interleukin-11 therapy selectively downregulates type I cytokine proinflammatory pathways in psoriasis lesions | Dec-99 | 12 | Completed |
| PMID: 24750799 | Immunomodulating treatment with low dose interleukin-4, interleukin-10 and interleukin-11 in psoriasis vulgaris | Jan-14 | 41 | Completed |
| **Mucositis** | | | | |
| NCT03720340 | Interleukin-11 Can Prevent and Treat of Radioactive Oral Mucitis | Aug-22 | 300 | Unknown status |

| | | | | |
|---|---|---|---|---|
| PMID: 35906582 | A prospective interventional study of recombinant human interleukin-11 mouthwash in chemotherapy-induced oral Mar-22 mucositis | Jul-22 | 112 | Completed |
| PMID: 11919725 | A phase I/II double-blind, placebo-controlled study of recombinant human interleukin-11 for mucositis and acute GVHD prevention in allogeneic stem cell transplantation | Mar-02 | 13 | Completed |
| **Other** | | | | |
| PMID: 26796134 | Four cases of investigational therapy with interleukin-11 against acute myocardial infarction | Jan-16 | 4 | Completed |
| PMID: 11438043 | Results of a phase-I/II randomized, masked, placebo-controlled trial of recombinant human interleukin-11 (rhIL-11) Jun-01 in the treatment of subjects with active rheumatoid arthritis | Apr-01 | 91 | Completed |
| PMID: 24613110 | Randomized clinical trial of human interleukin-11 in Dengue fever-associated thrombocytopenia | Mar-14 | 40 | Completed |

**Supplementary table 1. Clinical studies involving administration of rhIL11.**

| Name | Sequence | Primer Type |
| --- | --- | --- |
| *Il11ra1* Forward | CATTACCTACACATCCTTCCC | Transgene Forward |
| *Il11ra1* Reverse | CCTCCCACACTCAGATTAAGA | Transgene Reverse |
| *Myh6*-Cre Forward | ATGACAGACAGATCCCTCCTATCTCC | Transgene Forward |
| *Myh6*-Cre Reverse | CTCATCACTCGTTGCATCATCGAC | Transgene Reverse |
| IMR8744 | CAAATGTTGCTTGTCTGGTG | Internal Positive Control Forward |
| IMR8745 | GTCAGTCGAGTGCACAGTTT | Internal Positive Control Reverse |

**Supplementary table 2. Primers used for genotyping.**

| Primer/ Probe | Sequence |
|---|---|
| Forward | 5'-ACGTCCTGAAGTCTCCTGCCA-3' |
| Reverse | 5'-GTAAGGTAGCGGGTGGGCAA-3' |
| Probe | 5'-ACCGCTGACCTGGCCTGGACTCCA-3' |

**Supplementary table 3. Custom taqman assay for *Il11ra1* exon 4.**

**Supplementary table 4 List of significantly differentially expressed cardiomyocyte genes following rmIL11 injection.** Differentially expressed gene list from cardiomyocyte fraction of cells from single nuclei RNA sequencing of left ventricular tissue from male C57BL/6J mice 3 hours after injection with rmIL11 (200 mcg/kg) or saline (n=5). Genes ranked by logFC. (False discovery rate<0.05)

Attached separately

**Supplementary table 5. List of differentially regulated regions in mouse myocardial DNA following rmIL11 injection.** List of differentially regulated DNA regions from ATACseq analysis of mouse myocardial tissue 3 hours after rmIL11 injection (200 mcg/kg) compared to saline in wild type male C57BL/6J mice. (shrunken logFC +/-0.3, Adjusted p-value<0.1**).**

Attached separately

| Rank | Motif | P-value | log P-pvalue | % of Targets | % of Background | STD(Bg STD) | Best Match/Details |
|---|---|---|---|---|---|---|---|
| 1 | 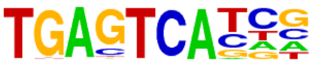 | 1e-447 | -1.031e+03 | 69.17% | 7.01% | 45.5bp (71.8bp) | FOSL2/MA0478.1/Jaspar(0.971) |
| 2 | 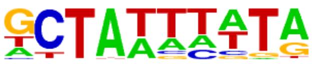 | 1e-44 | -1.035e+02 | 21.28% | 6.32% | 49.0bp (64.0bp) | Mef2d(MADS)/Retina-Mef2d-ChIP-Seq(GSE61391)/Homer(0.870) |
| 3 | 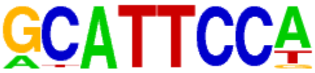 | 1e-22 | -5.112e+01 | 11.25% | 3.44% | 51.7bp (62.2bp) | TEAD1(TEAD)/HepG2-TEAD1-ChIP-Seq(Encode)/Homer(0.957) |
| 4 | 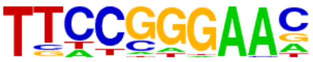 | 1e-20 | -4.703e+01 | 6.41% | 1.27% | 44.7bp (59.4bp) | Stat3(Stat)/mES-Stat3-ChIP-Seq(GSE11431)/Homer(0.950) |
| 5 | 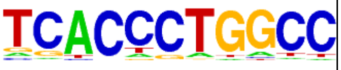 | 1e-15 | -3.531e+01 | 4.96% | 1.03% | 53.4bp (65.3bp) | ERE(NR),IR3/MCF7-ERa-ChIP-Seq(Unpublished)/Homer(0.685) |
| 6 | 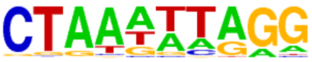 | 1e-14 | -3.446e+01 | 4.23% | 0.76% | 59.5bp (58.0bp) | Mef2b(MADS)/HEK293-Mef2b.V5-ChIP-Seq(GSE67450)/Homer(0.693) |
| 7 | 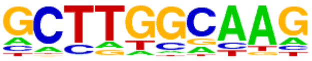 | 1e-14 | -3.255e+01 | 33.13% | 21.50% | 53.8bp (64.2bp) | NFIC/MA0161.2/Jaspar(0.818) |
| 8 | 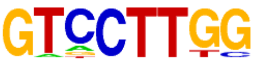 | 1e-13 | -3.068e+01 | 12.45% | 5.57% | 55.6bp (63.3bp) | SF1(NR)/H295R-Nr5a1-ChIP-Seq(GSE44220)/Homer(0.798) |

**Supplementary table 6. Enriched ATACseq transcription factor binding motifs.** Significantly positively enriched transcription factor binding motifs following de novo Homer analysis of ATAC-seq data from rmIL11 treated left ventricular tissue compared to saline controls (n=4).

| | Control + saline (n=4) | m6CMKO + saline (n=4) | Control + rmIL11 (n=4) | m6CMKO + rmIL11 (n=4) | p-value Control+IL11 vs m6CMKO+IL11 |
|---|---|---|---|---|---|
| **Male** | | | | | |
| **HR (bpm)** | 423 ± 3.5 | 492 ± 36 | 548 ± 14 | 434 ± 14.8 | 0.006 |
| **LVEF (%)** | 58.2 ±3.4 | 58.7 ±4.4 | 32.4 ±5.6 | 61.7 ±6.8 | 0.0004 |
| **VTI (mm)** | 44.7 ±2.9 | 35.4 ±2.6 | 25.4 ±1.9 | 34.1 ±1.4 | 0.045 |
| **GCS (%)** | -27.2 ±2.0 | -25.3 ±0.61 | -13.5 ±2.9 | -26.2 ±3.1 | 0.007 |
| **Female** | | | | | |
| **HR (bpm)** | 442 ±17 | 493 ±31 | 558 ±11 | 441 ±19 | 0.0005 |
| **LVEF (%)** | 64.9 ±2.5 | 62.8 ±3.9 | 32.5 ±2.1 | 61 ±2.6 | 0.0005 |
| **VTI (mm)** | 30.8 ±1.8 | 29.3 ±3.3 | 18.2 ±1.2 | 35 ±0.29 | 0.0001 |
| **GCS (%)** | -29.6 ±2.6 | -30.7 ±2.3 | -11.5 ±1.2 | -28.5 ±2.4 | 0.0002 |

**Suppl table 7. Echocardiographic measurements in IL11 treated m6CMKO mice.** Genetic cardiomyocyte-specific IL11 receptor knockout mice (m6CMKO) and *Il11ra1*$^{fl/fl}$ controls (Control) were treated with saline (2 uL/kg) or rmIL11 (200 mcg/kg) for 2 hours. Echocardiographic parameters were recorded under isoflurane anaesthetic. Values are reported as mean±standard error of the mean. *Statistics: Comparison between groups by one-way ANOVA with Sidak's multiple comparisons. P-values less than 0.05 are considered significant.* Abbreviations used **HR**, heart rate; **bpm**:, beats per minute, **LVEF**, left ventricular ejection fraction; **VT**I, velocity time integral from pulse wave doppler trace in the aortic arch; **GCS**, global circumferential strain.

#### Supplementary Figures

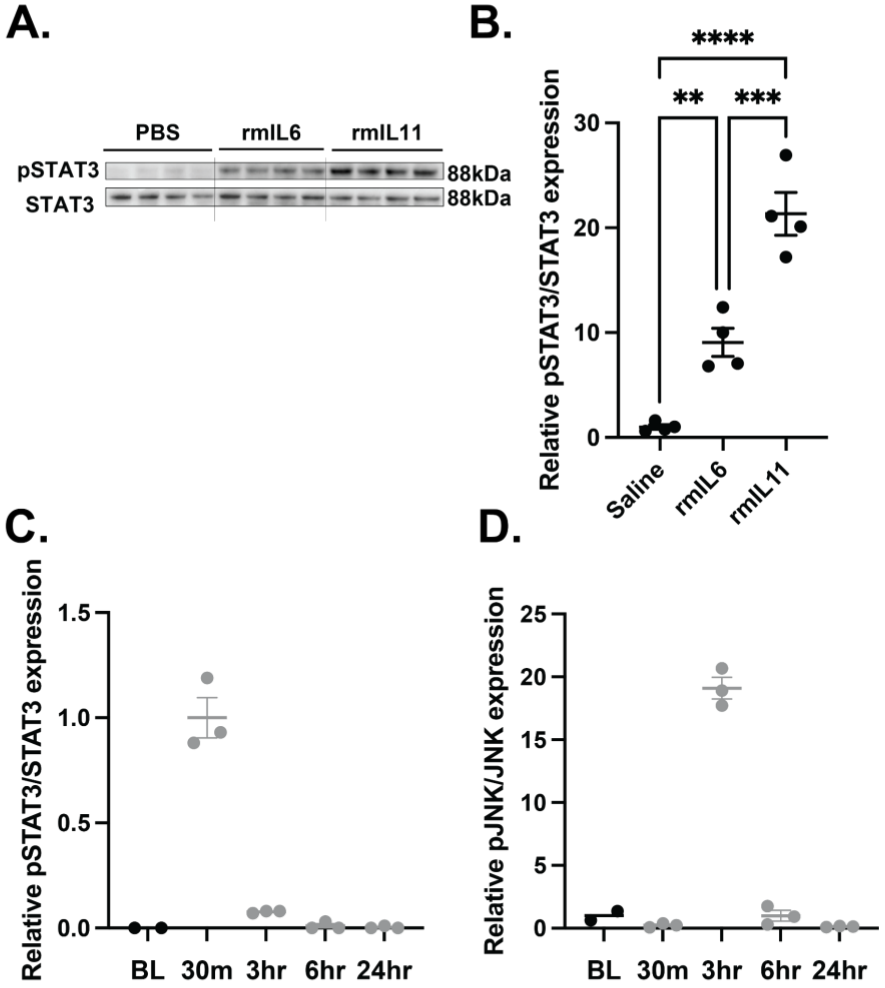


**Supplementary Figure 1. IL11 and IL6 signalling within the heart. (A)** Western blot of pSTAT3/STAT3 from male C57BL/6J mice treated with 200 mcg/kg of rmIL6, rmIL11 or an equivalent volume of saline. **(B)** Quantification of western blot from (A). Quantification of expression of **(C)** pSTAT3 compared to total STAT3 and **(D)** pJNK compared to total JNK from Figure 1i. *Statistics: One-way ANOVA with Sidak's multiple comparison testing. Significance denoted as **p<0.01, ***p<0.001, ****p<0.0001*

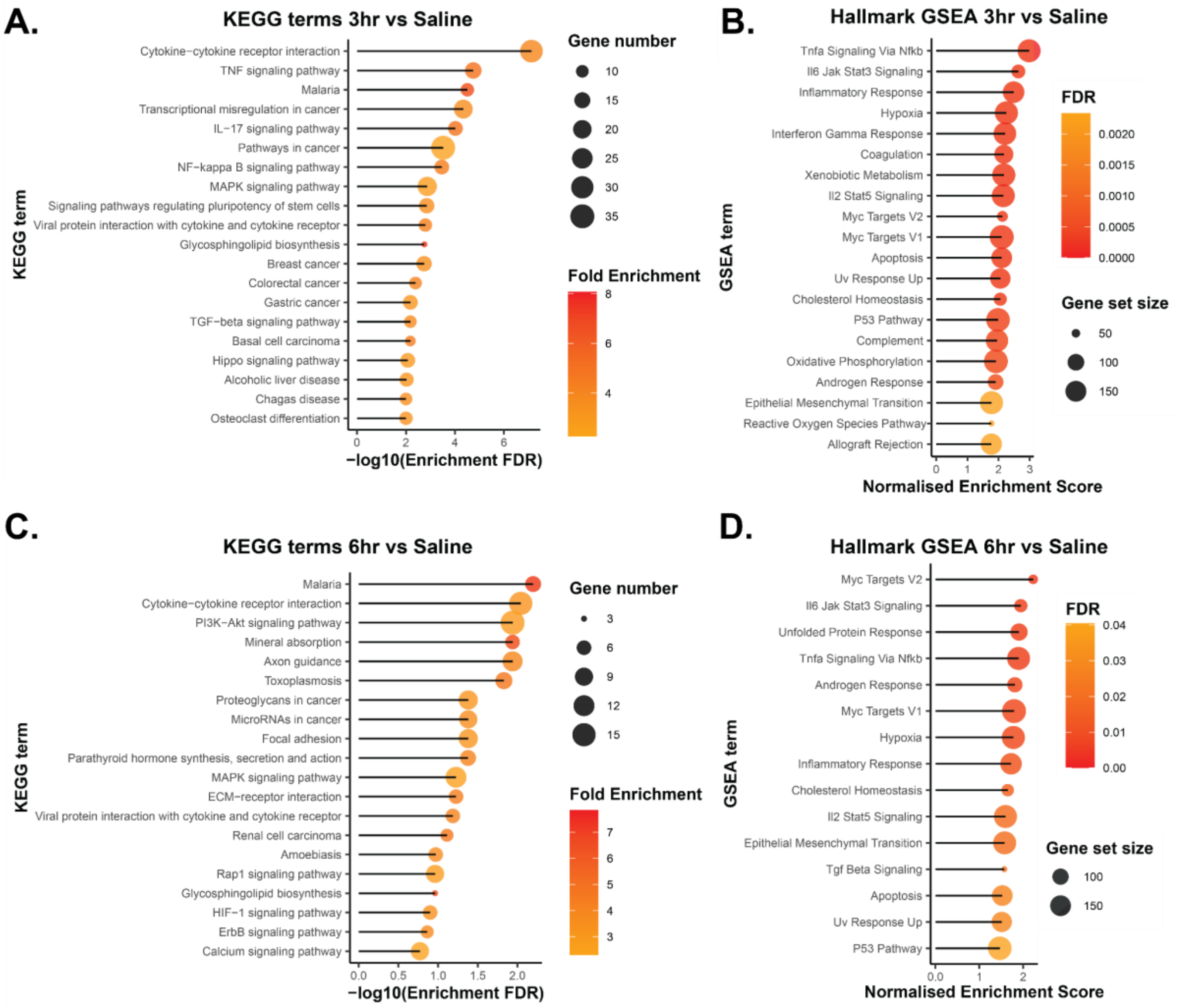


**Supplementary Figure 2. Bulk myocardial RNA-sequencing at 3 and 6 hours.** Analysis of differentially expressed genes from left ventricle RNA seq data following rmIL11 injection compared to saline injection. **(A)** Top 20 most significantly enriched KEGG terms 3 hours after injection (n=4) and **(B)** top 20 most enriched GSEA terms 3 hours after injection (n=4). **(C)** Top 20 most significantly enriched KEGG terms 6 hours after injection (n=4) and **(D)** all significantly enriched GSEA terms 6 hours after injection (n=4).

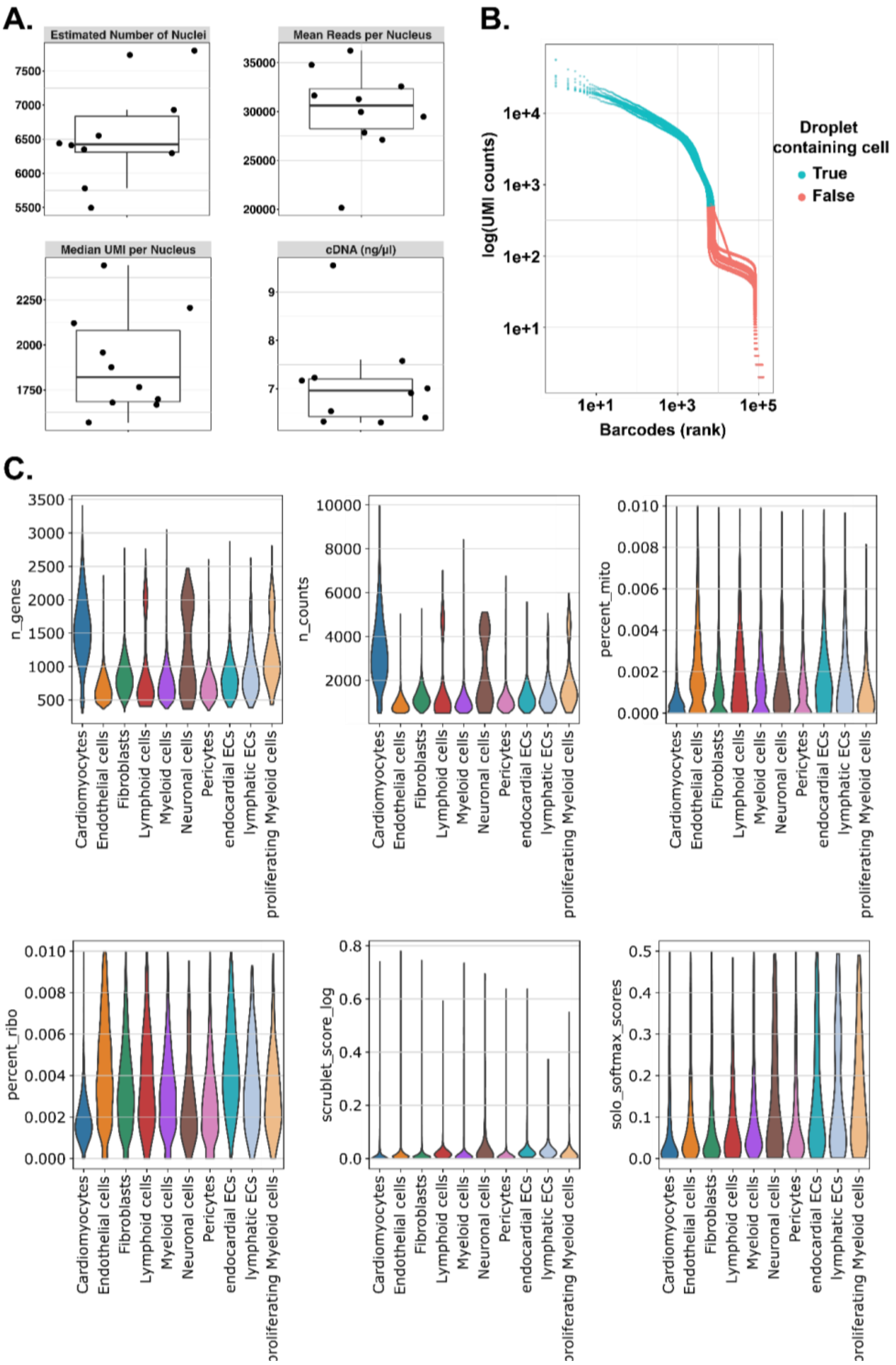


**Supplementary Figure 3. Quality assessment of snRNAseq data. (A)** Estimated number of nuclei, mean reads and median unique molecular identifier (UMI) count per nuclei, and cDNA concentration (ng/µl). **(B)** Colour coded barcode rank plots across all samples shows classification of nuclei. Turquoise represents nuclei-containing droplets (true); Red represents empty droplet, background, or ambient RNA (false). A clear distinction between nuclei containing droplets and empty droplets (background ambient RNA) indicates a low overall background. **(C)** Violin plots show number of genes (n_genes), number of UMIs (n_counts), percent UMIs mapping to mitochondrial (percent_mito) and ribosomal genes (percent_ribo) and doublet scores (scrublet_score / solo_score) plotted per cell type after quality control filtering and clustering.

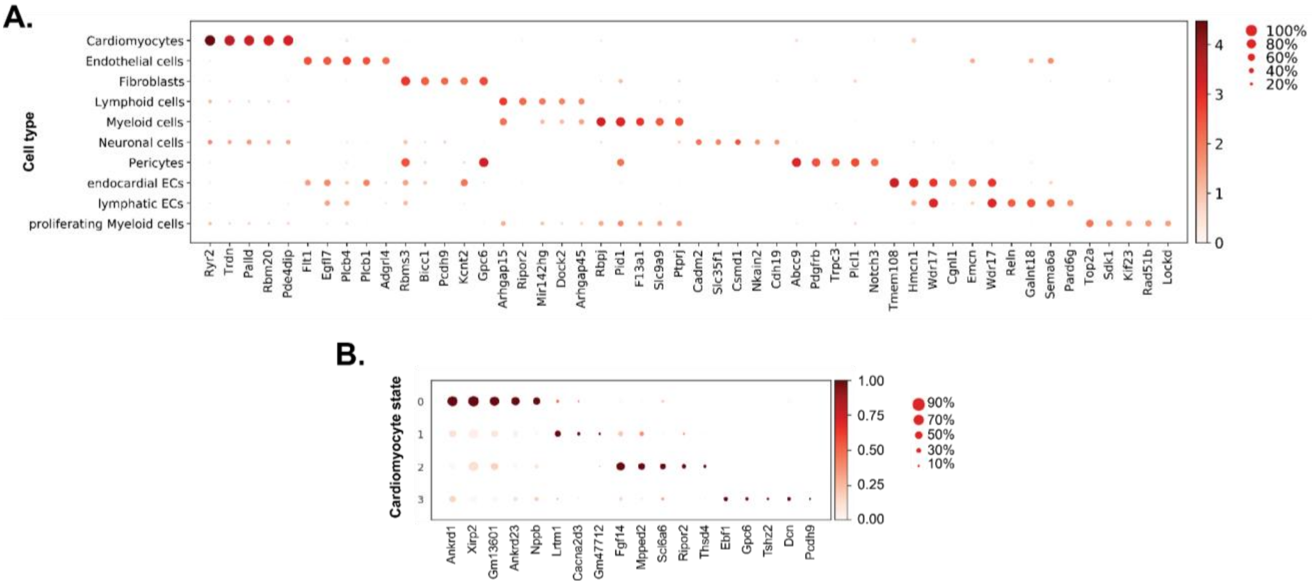


**Supplementary figure 4. SnRNAseq marker genes. (A)** Dotplot showing selected marker genes of each cell type. **(B)** Dot plot showing selected marker genes for each cardiomyocyte cell state. Dot size represents fraction (%) of expressing cells and colour represents mean expression level.

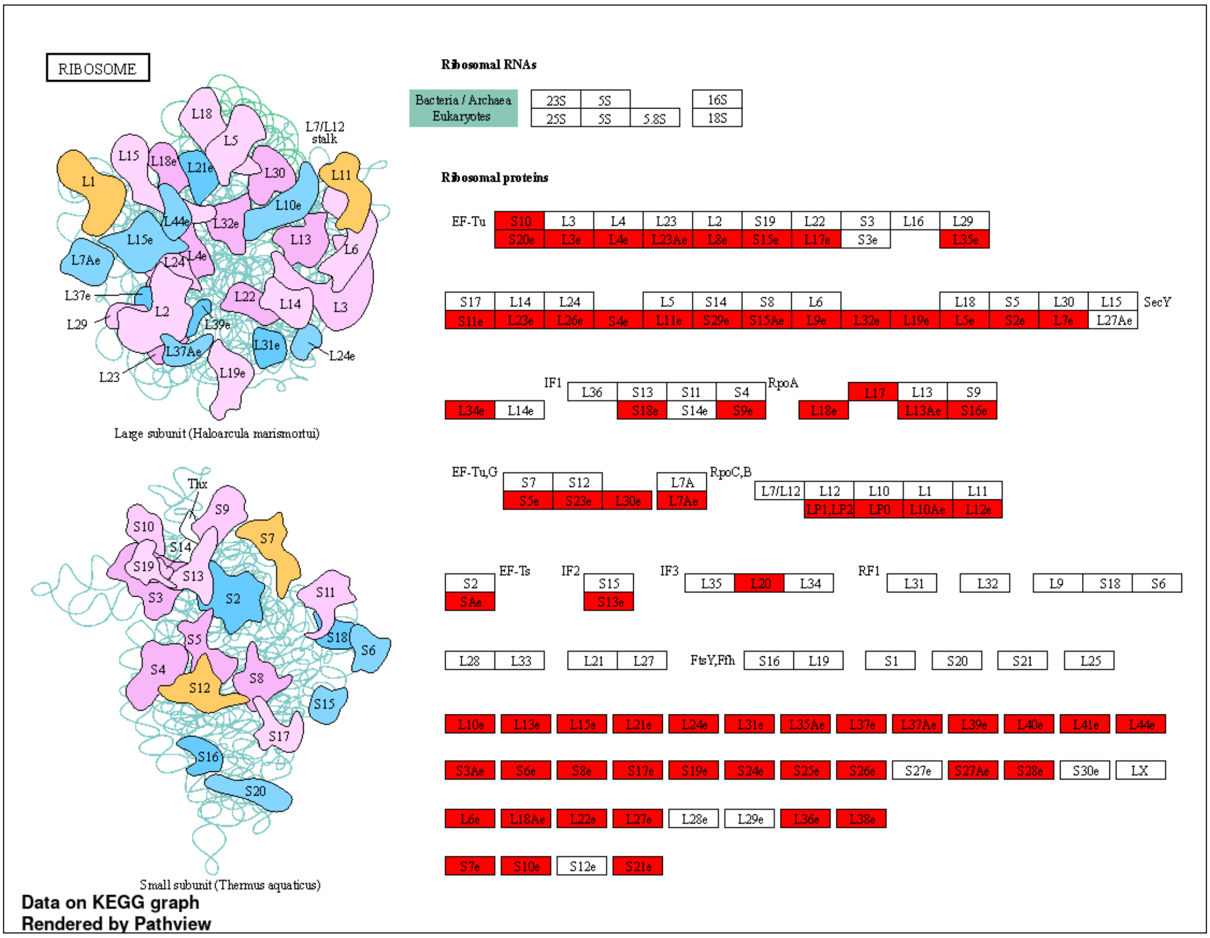


**Supplementary Figure 5. Ribosome KEGG analysis.** Illustration of the most significantly enriched KEGG pathway from snRNAseq analysis following administration of rmIL11. Significantly differentially expressed genes in cardiomyocytes from snRNAseq analysis are highlighted in red.
