## Supplementary figures and images for "Interleukin 11 therapy causes acute heart failure and its use in patients should be reconsidered"

### Uncropped blots

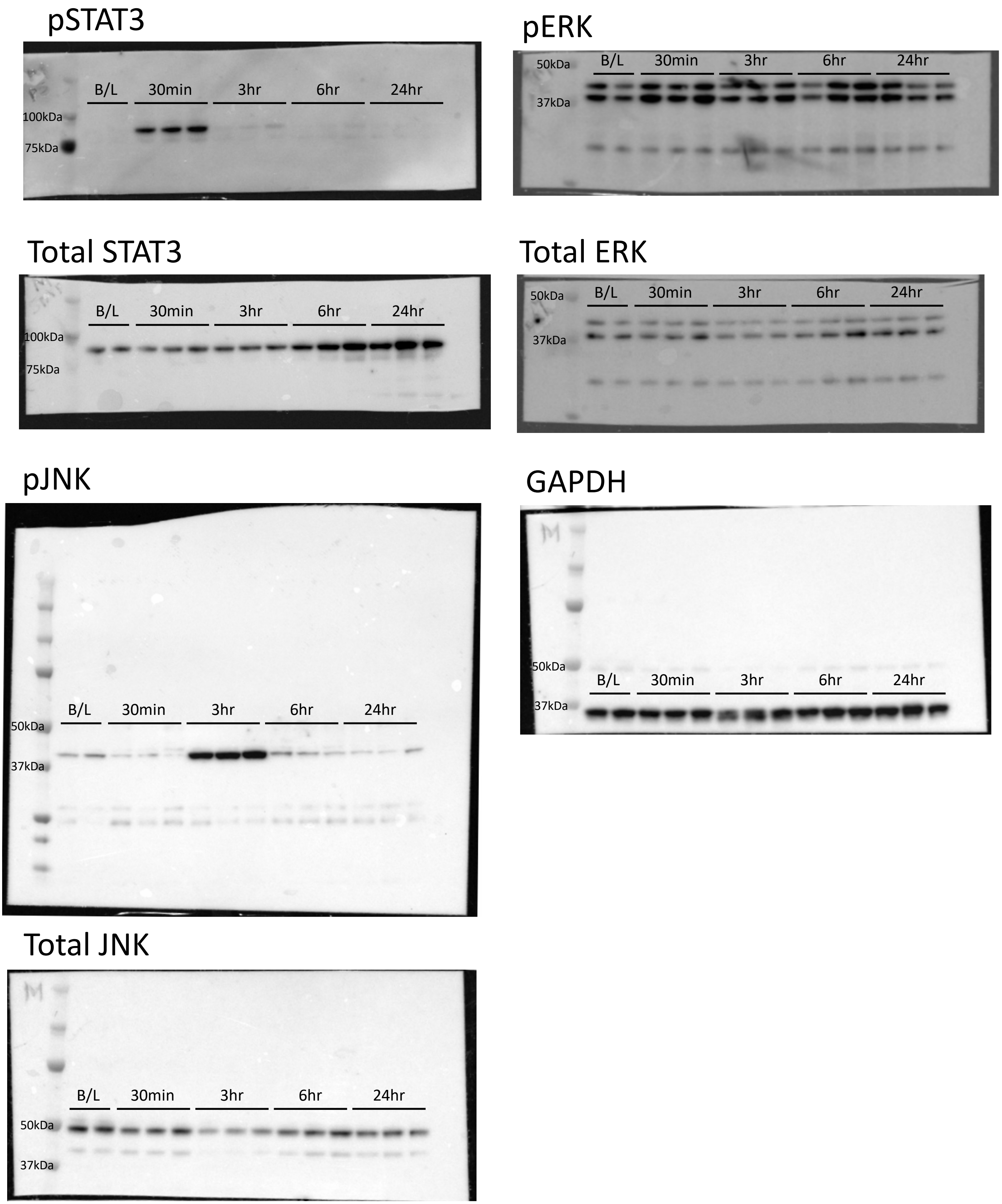

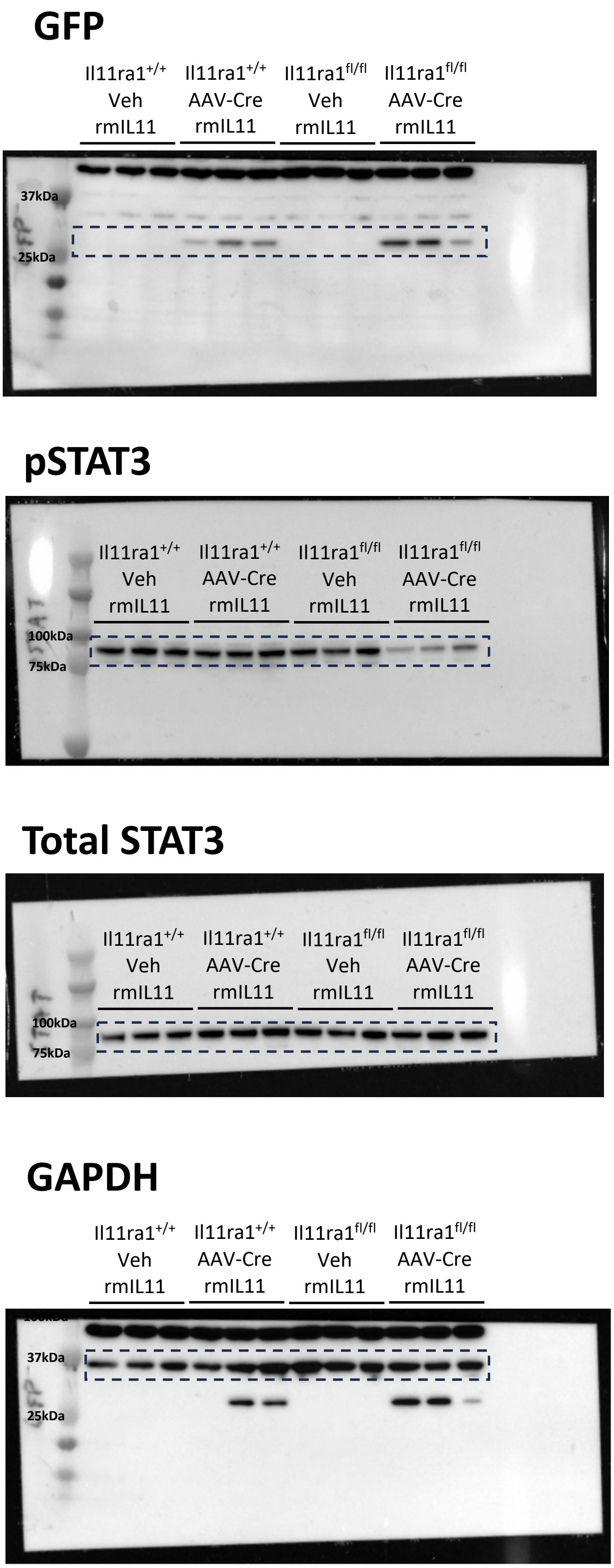

#### Figure 6C

##### Males

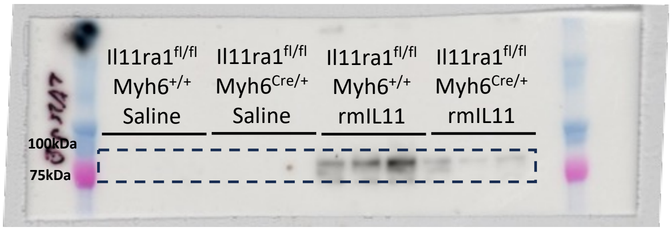


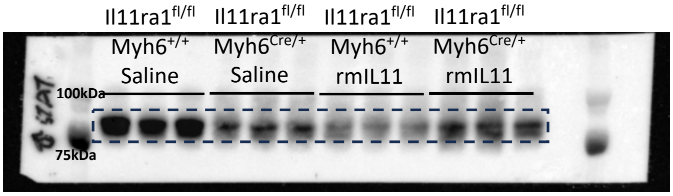


##### Females

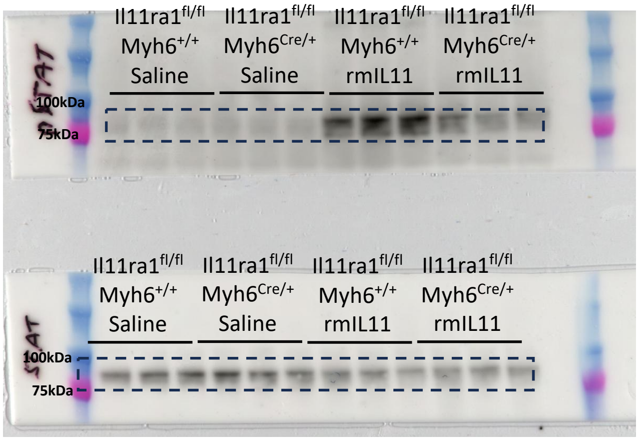

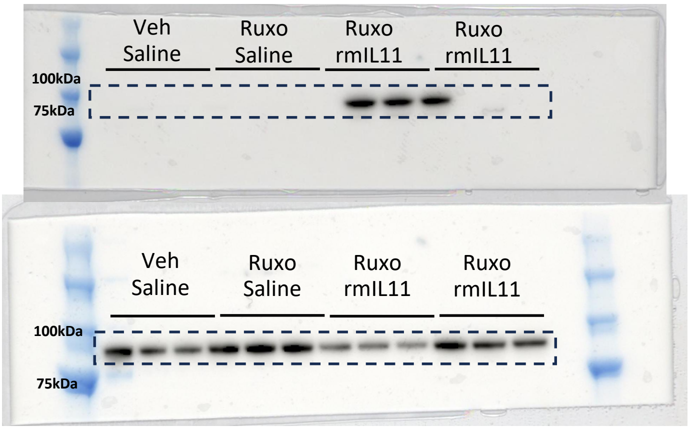
